# Dopamine enhances multisensory responses in the dorsomedial striatum

**DOI:** 10.1101/2023.09.25.559270

**Authors:** María Sáez, Javier Alegre-Cortés, Nicolás A. Morgenstern, Cristina García-Frigola, Roberto de la Torre-Martínez, Ramón Reig

## Abstract

The brain operates with simultaneous different sensory modalities in order to engage adaptive responses. However, the question of how (and where) multisensory information is integrated remains unanswered. In the dorsomedial striatum, single medium spiny neurons (MSNs) are excited by tactile and visual inputs; however, the mechanism which allows the integration of these responses and how they are shaped by dopamine is unknown.

Using *in vivo* optopatch-clamp recordings, we study how dopamine modulates tactile, visual and simultaneous bimodal responses in identified MSNs and their spontaneous activity. Results show that dopamine enhances bimodal responses, specifically in direct pathway MSNs, through the acceleration of the visual responses. We provide anatomical and computational evidence suggesting that this relies on the disinhibition of direct MSNs by a cell-type-specific corticostriatal pathway. Altogether, our *in vivo*, *in silico* and tracing results propose a new mechanism underlying the synchronization of multimodal information mediated by dopamine.

## INTRODUCTION

The brain is constantly processing information conveyed by several sensory modalities to create accurate representations of the world, allowing adaptive behaviors in a changing environment. Most of our motor activities require, at least, the combination of visual and somatosensory information to succeed. Both sensory modalities are used as feedback signals during actions^1^. However, the mechanisms underlying how visual and tactile information are combined in the brain to produce accurate responses are not fully understood.

The basal ganglia (BG) play an essential role in motor control^2,3^, sensorimotor functions and motor learning^4,5^, as well as decision making and reward^6,7^, all of which require the integration of sensory information. Malfunctions of BG nuclei lead to severe disorders such as Parkinson’s disease among others^8^, where motor symptoms coexist with several sensory problems that emerge at early stages of the disorder^9,10^.

The striatum, the input nucleus of the BG, receives cortical projections from sensory, motor and associative areas as well as from the thalamus^11–14^. In rodents, the dorsal striatum is divided into two different functional regions^15,16^: the dorsolateral (DLS) and the dorsomedial (DMS) striatum, both participating in motor control. However, each of them receives specific sets of cortical projections from intratelencephalic (IT) and pyramidal tract (PT) neurons^17,18^. Axons from different cortical sensory areas target striatal GABAergic projection medium spiny neurons from both the direct (dMSNs) and indirect pathway (iMSNs), as well as interneurons^19–21^. A critical regulator of striatal microcircuits are cholinergic interneurons (ChINs)^22^, characterized by tonic discharge, which reduces the excitability of MSNs by disynaptic inhibition^23–25^.

The dorsal striatum is also densely innervated by dopaminergic axons from the *Substantia nigra pars compacta* (SNc), which interact with cholinergic interneurons ^26–29^, modulating corticostriatal transmission and plasticity *in vivo*^30^ and behaviorally having been linked to a great variety of functions related with motor control, reinforcement and expectation^31^. The release of DA in the striatum impacts on dMSNs and iMSNs activity via the activation of D1 or D2 receptors, respectively^32–37^. Furthermore, the activation of DA neurons correlates with the pauses of tonic discharge of ChINs, *ex vivo*^38–40^ and *in vivo*^41^, promoting in turn, the disinhibition of MSNs activity^42,43^.

In the DMS, both MSNs types respond to visual and tactile stimulation^44^. These responses are characterized by particular latencies in which visual cues are processed tens of milliseconds slower than tactile ones, mainly attributed to retinal processing^45^. This result in the independent synaptic integration of tactile and visual inputs^44^. However, how these inputs from different sensory modalities and temporal discrepancies are integrated remains underexplored.

Here, using *in vivo* optopatch-clamp recordings, we measured the activity of identified dMSNs and iMSNs in the mouse DMS, responding to visual, whisker or simultaneous bimodal stimulation in resting conditions or after optogenetic DA release. Our data reveals that DA enhances the integration of bimodal responses in dMSNs by synchronizing visual and tactile inputs, supported by a specific corticostriatal projection from primary visual cortex to the DMS. This idea was supported by the construction of a computational model following anatomical constrains which reproduced and explained the results obtained *in vivo*; suggesting that multisensory synchronization on dMSNs is induced by the pause of ChINs and DA release. These results represent a step forward towards understanding the role of DA in the striatal microcircuits and decipher a new mechanism by which the brain is able to integrate information from different sensory modalities.

## RESULTS

### *In vivo* whole-cell optopatch-clamp recordings in dorsomedial MSNs

MSNs are characterized by low frequency of action potentials^46–48^ and most of their synaptic activity results in subthreshold depolarization. Thus, in order to understand synaptic integration on MSNs, we acquired *in vivo* whole-cell patch-clamp recordings, allowing the study of subthreshold voltage dynamics on MSNs. We recorded 38 striatal neurons of the DMS in D2-cre x ChR2 x DAT-cre mice (n=12), under ketamine and medetomidine anaesthesia (Fig. 1A-C). The use of anaesthesia enables the activation of sensory pathways while avoiding interference with motor-related inputs. Under these conditions, the membrane potential of MSNs exhibits a bimodal distribution known as Slow Wave Oscillation (SWO), switching from Down to Up states^49^. All the recorded neurons displayed prominent SWO (Fig. 1D, E; 2A; 3A), as described before^15,44,49,50^. From the 38 recorded neurons, 33 were classified as MSNs, 2 as Fast Spiking Interneurons (FSIs) and 3 as ChINs (Fig. Supp. 1A; Table 1). MSNs were identified from other striatal types by their electrophysiological properties as well as their morphology and optogenetic identification (Fig 1B, D, G-I). FSIs and ChINs were distinguishable by their electrophysiological properties (Table 1). Simultaneously, local field potential (LFP) recordings were obtained from S1 and V1 to ensure that visual and tactile stimulations elicited responses in their respective cortical areas (Fig. 1A, D; 3A). MSNs from the direct and indirect pathways were identified using the optopatcher^15,51^ (Fig. 1E). Depolarizing neurons were categorized as iMSNs, whereas non-depolarizing MSNs were classified as putative dMSNs. Input resistance was measured in both MSNs types for Up and Down states. Depolarization increased the input resistance of MSNs, either induced by current injection (Fig. 1F) or by the spontaneous Up states (Table 1, Fig. Supp. 2). Our results indicate similar resistance, capacitance, tau (Fig. 1G-I), membrane potential and SWO frequency values when comparing both types of MSNs (Table 1). Therefore, we conclude that both subpopulations display closer electrophysiological properties in the DMS, in contrast with dMSNs and iMSNs of the DLS^44,51,52^. This similitude in the DMS support particular functional properties between pathways in the DLS and DMS, as shown previously^15^.

**Figure 1.**
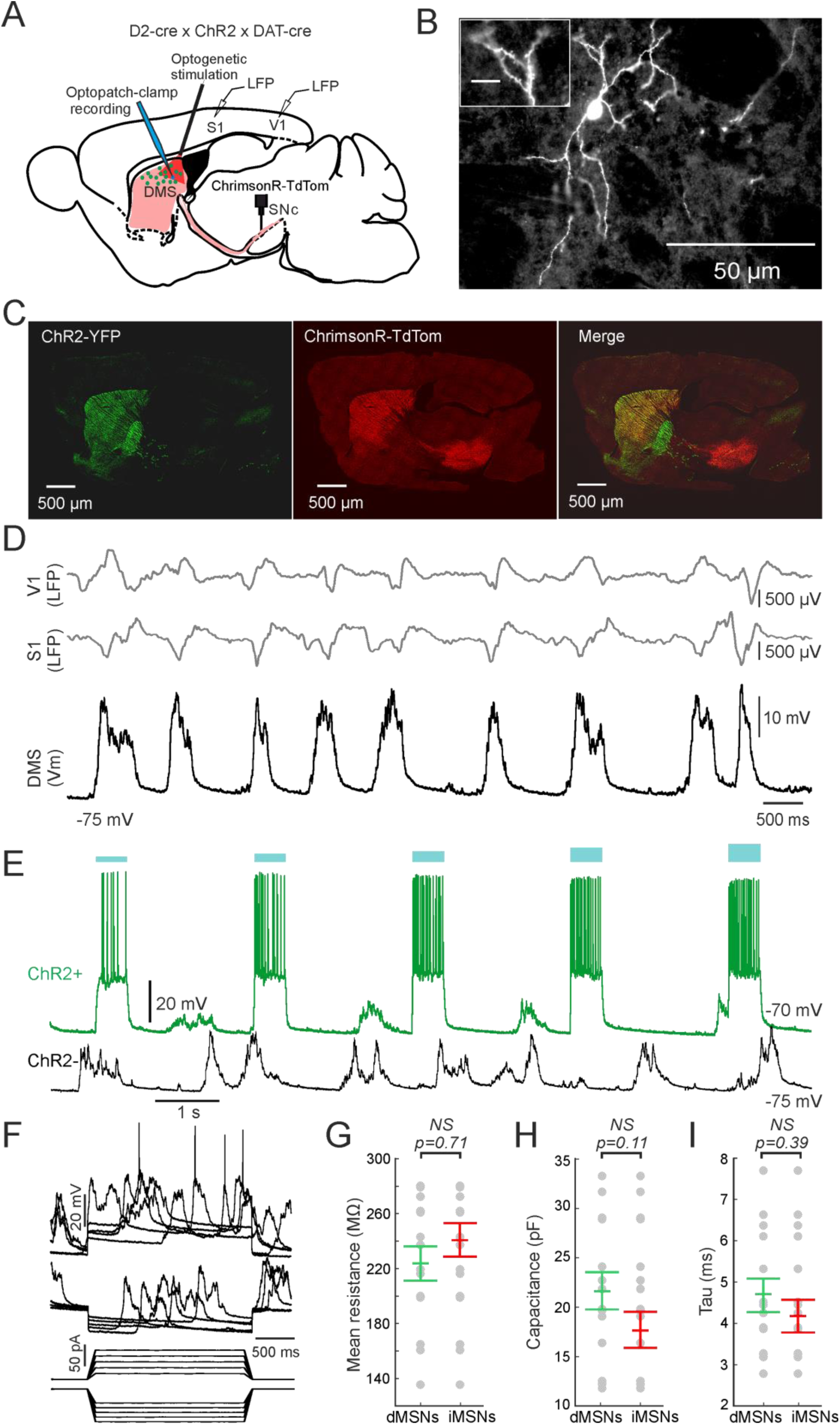
Experimental approach. **A,** Schematic representation of the experimental set up, in vivo whole-cell optopatch-clamp recordings with simultaneous LFP in S1 and V1. **B,** Morphological reconstruction of a dorsomedial MSN. Different scales show neuron magnitude and its dendritic spines, confirming that the recorded neuron is a MSN. Inset bar scale corresponds to 20 µm. **C,** Sagittal images of a D2-cre x ChR2 x DAT-cre mice injected with the virus expressing the opsin ChrimsonR-TdTomato in the SNc. Left: Expression of ChR2-YFP in the iMSNs. Middle: Dopaminergic terminals of the SNc expressing the opsin ChrimsonR-TdTomato in the SNc after at least 8 weeks post injection. Right: Merge of both images. Notice that the yellow colour indicates the merge between the dopaminergic terminals and the iMSNs expression only in the striatum. **D,** Example of a recorded MSN in the DMS (black trace) with simultaneous LFP in S1 and V1 (grey traces). **E,** Example showing an in vivo identification of a MSN using the optopatcher. Indirect MSNs in the D2-cre x ChR2 x DAT-cre mice (top trace, ChR2+, green) responded to light pulses, inducing a depolarization in the MSN. Negative cells (bottom trace, ChR2-, black) did not respond to light pulses. Blue bars indicate the intensity of the light pulse stimulation, from 20 to 100 % (0.166 mW and 0.83 mW, respectively). **F,** Response of an MSN to hyperpolarizing step current injections. **G, H, I,** Mean resistance (G), Capacitance (H) and Tau (I) values calculated for dMSNs and iMSNs.

**Table 1.** Intrinsic properties of dorsomedial dMSNs and iMSNs, ChINs and FS. . All values are means ± standard deviation. MSNs: n=33 (18 dMSNs; 15 iMSNs). ChINs: n=3. FS: n=2.

|  | All MSNs | dMSNs | iMSNs | ChINs | FS |
| --- | --- | --- | --- | --- | --- |
| Input Resistance (MΩ) | 232 ± 54.74 | 223 ± 46.82 | 240 ± 61.69 | 203 ± 37.87 | 250 ± 21.92 |
| Resistance Down state hyp. (MΩ) | 211 ± 60.47 | 201 ± 43.62 | 221 ± 72.99 | 205 ± 44.37 | 239 ± 20.50 |
| Resistance Down state dep. (MΩ) | 240 ± 65.28 | 238 ± 72.90 | 241 ± 59.88 | 186 ± 55.56 | 263 ± 25.45 |
| Resistance Up state hyp. (MΩ) | 240 ± 61.62 | 220 ± 58.11 | 259 ± 60.70 | 208 ± 45.61 | 257 ± 22.62 |
| Resistance Up state dep. (MΩ) | 254 ± 70.24 | 245 ± 63.58 | 262 ± 77.23 | 185 ± 48.41 | 224 ± 11.31 |
| Capacitance (pF) | 19.59 ± 6.23 | 21.65 ± 7.13 | 17.67 ± 4.71 | 32.38 ± 24.1 | 16 ± 7.1 |
| Tau (ms) | 4.43 ± 1.42 | 4.69 ± 1.50 | 4.19 ± 1.34 | 6.09 ± 3.38 | 3.52 ± 0.76 |
| Sag (mV) | 0 ± 0 | 0 ± 0 | 0 ± 0 | 7.78 ± 2.20 | 0 ± 0 |
| Membrane potential (mV) | -68.62 ± 5.49 | -69.26 ± 5.37 | -68.04 ± 4.11 | -51 ± 6.55 | -65 ± 7.07 |
| Firing rate (Hz) | 0.197 ± 0.39 | 0.182 ± 0.32 | 0.211 ± 0.43 | 1.35 ± 0.41 | 2.69 ± 0.95 |
| Oscillation frequency (Hz) | 0.69 ± 0.06 | 0.68 ± 0.06 | 0.71 ± 0.07 | 0.79 ± 0.08 | 0.72 ± 0.01 |

### Dopamine decreases the spontaneous high frequency oscillations during Up states

In order to explore the impact of DA on the spontaneous activity of MSNs, we stimulated optogenetically dopaminergic axons in the DMS. Before, fiber photometry experiments were performed^53–55^ to confirm the release of DA. Briefly, optogenetic stimulation of dopaminergic axons (4 pulses, each of them of 15 ms at 15 Hz), produced a reliable DA release in the DMS (Fig. Supp. 3).

During the Up states of the SWO, several bands of high frequency oscillatory activity can be recorded in MSNs^15^ as well as in the cortex^56,57^, thus, we computed the energy of the theta- (6- 10 Hz), beta- (10-20 Hz) and gamma-bands (20-80 Hz) [see methods] to assess the impact of DA. Our results showed that DA decreased the energy of all of them in both MSNs subpopulations: theta- (control= 2.20 ± 0.63 mV, DA= 1.79 ± 0.45 mV2, p=0.000003); beta- (control= 3.11 ± 0.55 mV2, DA= 2.72 ± 0.41 mV2, p=0.00009); and gamma-band (control= 0.60 ± 0.33 mV, DA= 0.36 ± 0.21 mV2, p=0.000004) (Fig. 2A, B (green and red traces)). This energy decrease was always accompanied with the TdTomato virus reporter expression in the DMS (Fig. 1C). To ensure that it was related to DA, we performed a set of control experiments. As described previously, mice were injected in the SNc with the cre-dependent control virus AAV5-CAG-FLEX-tdTomato. In this case, optogenetic stimulation did not modify the energy in any of the explored frequency bands (Fig. 2B, black and grey traces), confirming that changes were induced by DA. When comparing dMSNs and iMSNs, differences were not detected (Fig. 2B).

**Figure 2.**
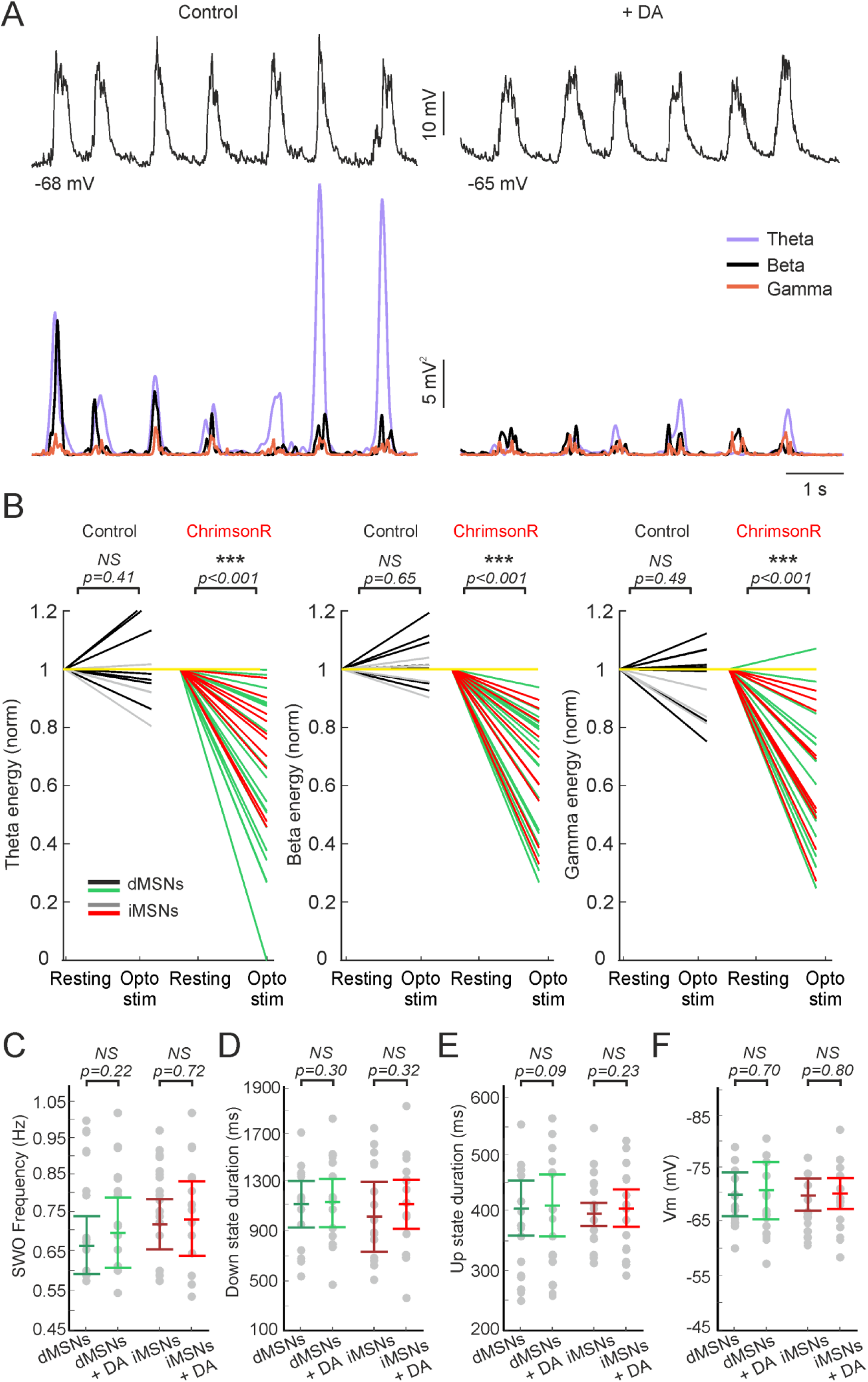
Dopamine modulation of spontaneous activity. **A,** Example traces of the SWO in MSNs in control (left upper trace) and during DA release (right upper trace). Coloured traces in the bottom part correspond to the theta-, beta- and gamma-band energy from each upper trace. Notice how the energy of all bands decreases when DA is released. **B,** Theta- (left), beta- (middle) and gamma-band (right) normalized energy of dMSNs and iMSNs. Black and grey lines correspond to photostimulated dMSNs and iMSNs expressing an empty construct (only tdTomato), respectively; while green and red lines correspond to photostimulated dMSNs and iMSNs expressing the opsin construct (ChrimsonR- TdTomato), respectively. Yellow line indicates the normalized value 1. Values were normalized to the average value presented in control conditions in each frequency band for each neuron. **C, D, E, F,** Quantification of the SWO frequency **(C)**, Down state duration **(D)**, Up state duration **(E)** and Vm (Voltage membrane potential) **(F)** of dMSNs and iMSNs DMS-MSNs in resting DA levels or after optogenetic stimulation.

In addition, we also observed that the SWO frequency, Up and Down state duration, as well as membrane voltage during Down state were not affected by DA (Fig. 2C-F). This suggests that local changes of DA do not modulate the slow oscillatory activity, although they do modulate the high frequency oscillations that occur during the up states.

Finally, several *ex vivo* studies described how dopaminergic neurons can co-release GABA ^58,59^ or glutamate ^60–64^. However, in our DMS *in vivo* recordings we did not detect any evident fast depolarization (EPSPs) or hyperpolarization (IPSPs) induced either by optogenetic or electrical stimulation of the SNc (Fig. Supp. 4A, B); even when injecting positive current, moving their membrane potential close to the excitatory reversal potential in order to study the inhibitory components (IPSPs) in some MSNs (n=6).

### Dopamine modulates bimodal responses specifically in dMSNs

We next tested the modulatory effects of DA on the integration of visual and tactile information by MSNs. We recorded 29 MSNs: 14 dMSN and 15 iMSNs, from D2-cre x ChR2 x DAT-cre mice (n=12) (Fig. 3A). Only recordings of MSNs with the whole sequence of stimulation were included, where the average recording time was 49.96 ± 10.27 minutes. We delivered single contralateral air puff stimulation to the whisker pad or a flash of white light, both with a duration of 15 milliseconds, every 5 seconds. In addition, we also stimulated with both sensory modalities simultaneously (bimodal responses). DA was released by optogenetic stimulation of dopaminergic terminals in the DMS, 30 milliseconds prior to sensory stimulation (Fig. 1A, C). Thus, tactile, visual and bimodal stimulation at resting or after DA release was randomly launched, evoking responses in all the recorded MSNs (Fig. 3A, B).

**Figure 3.**
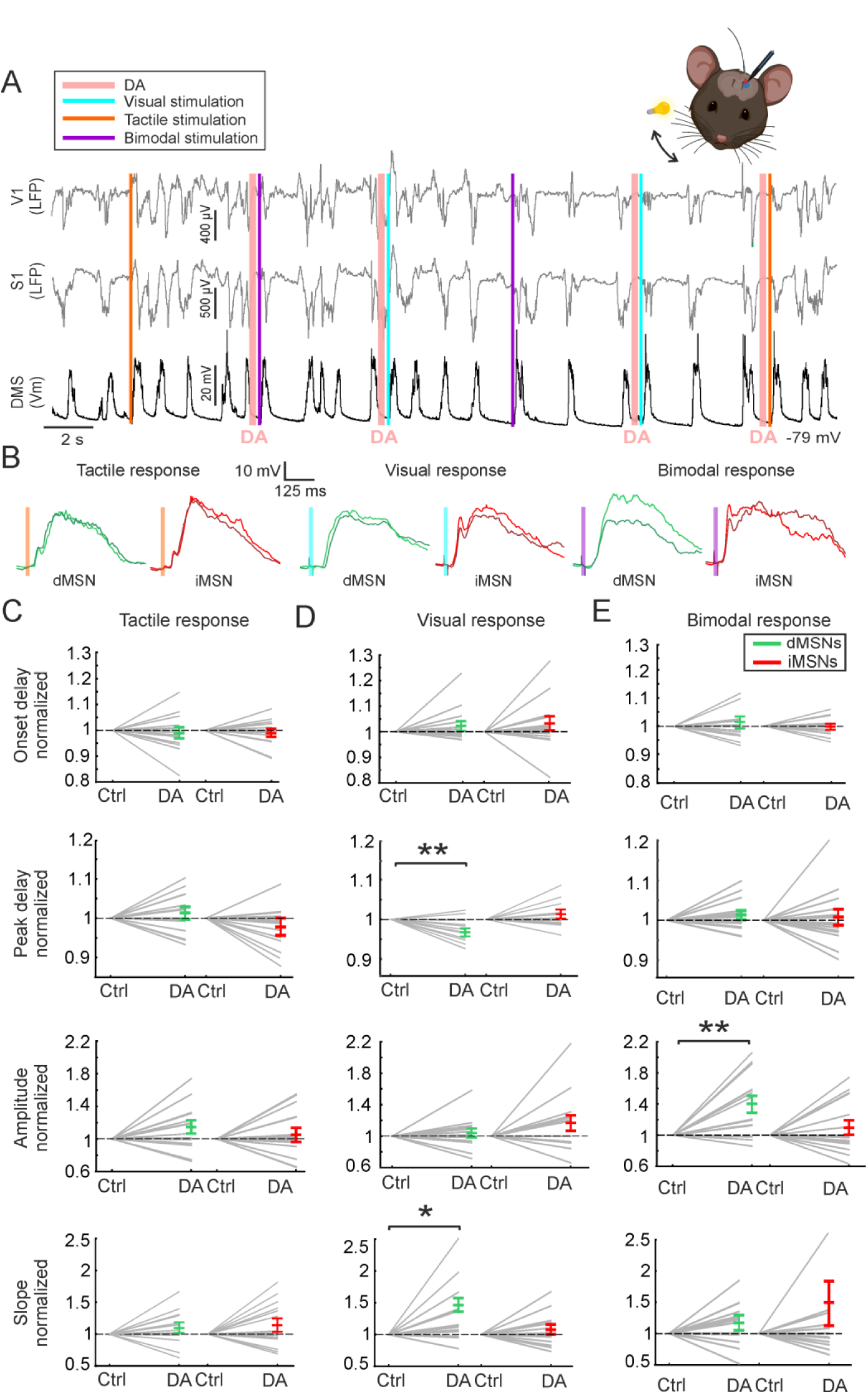
Dopaminergic modulation of sensory responses in DMS. **A,** Example of an in vivo whole-cell patch-clamp recording (black trace) with simultaneous LFPs in V1 and S1 (grey traces), while performing optogenetic, visual, tactile and bimodal stimulation randomly. **B,** Waveform averages of dMSNs (green) and iMSNs (red) MSNs for the different sensory stimulations in presence of DA (light colours) or control conditions (dark colours). Vertical coloured line indicates the time of the sensory stimulation. **C, D, E,** Normalized averages of dMSNs and iMSNs responses after DA release, for tactile **(C)**, visual **(D)** and bimodal **(E)** stimulation. The normalization was done with respect to the average value presented in control conditions during the Down states for each neuron. * p<0.05, ** p<0.01. Raw data can be found in Table 2.

**Table 2.**
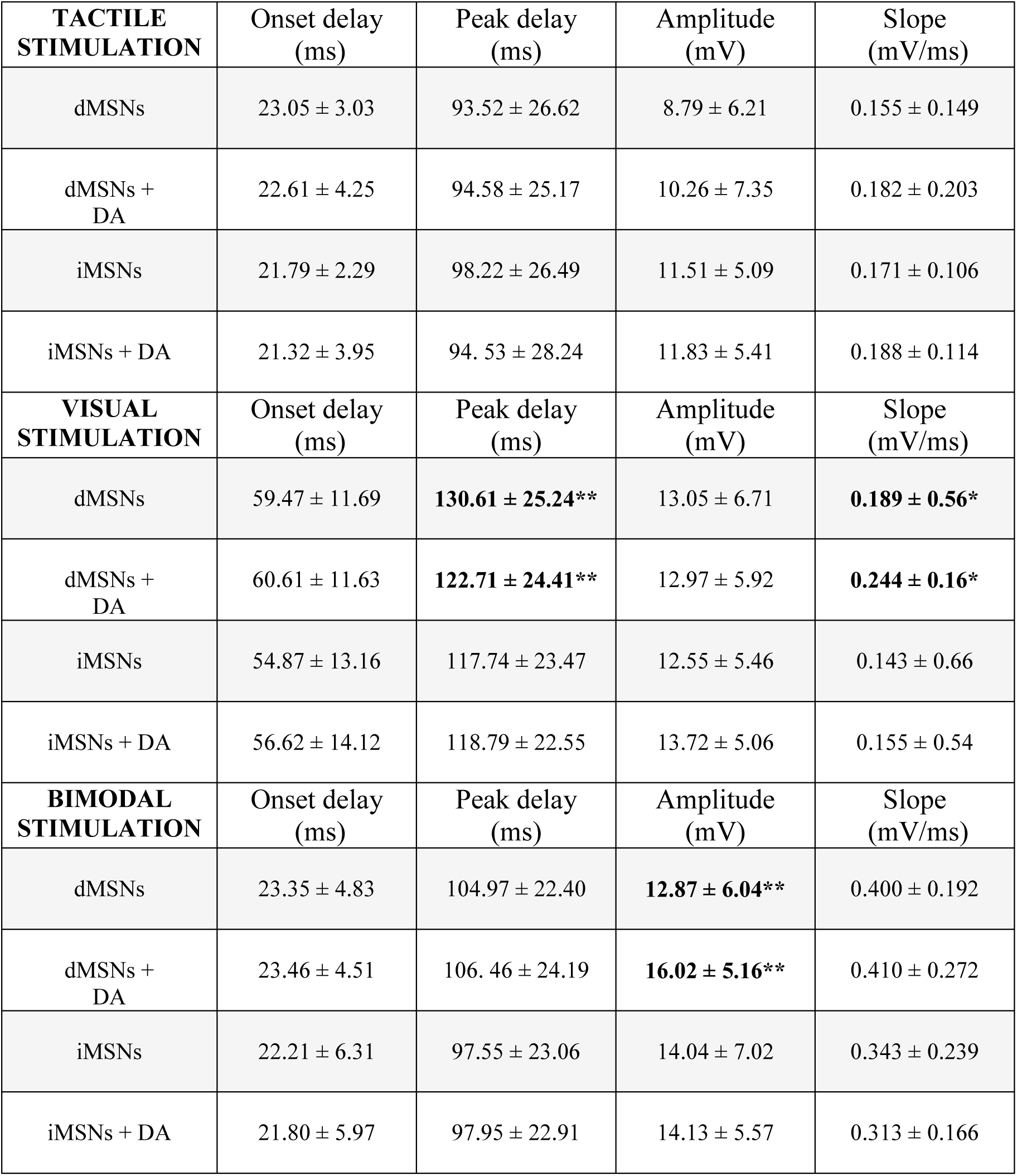
Mean values of tactile, visual and bimodal responses for direct and indirect MSNs with or without DA release. All values are mean ± standard deviation. n=29 (14 dMSNs; 15 iMSNs). Asterisks indicate significant differences when comparing control and photostimulated dMSNs and iMSNs. * p<0.05, ** p<0.01. No differences were observed when comparing dMSNs and iMSNs.

| **TACTILE STIMULATION** | Onset delay (ms) | Peak delay (ms) | Amplitude (mV) | Slope (mV/ms) |
| --- | --- | --- | --- | --- |
| dMSNs | 23.05 ± 3.03 | 93.52 ± 26.62 | 8.79 ± 6.21 | 0.155 ± 0.149 |
| dMSNs + DA | 22.61 ± 4.25 | 94.58 ± 25.17 | 10.26 ± 7.35 | 0.182 ± 0.203 |
| iMSNs | 21.79 ± 2.29 | 98.22 ± 26.49 | 11.51 ± 5.09 | 0.171 ± 0.106 |
| iMSNs + DA | 21.32 ± 3.95 | 94.53 ± 28.24 | 11.83 ± 5.41 | 0.188 ± 0.114 |
| **VISUAL STIMULATION** | Onset delay (ms) | Peak delay (ms) | Amplitude (mV) | Slope (mV/ms) |
| dMSNs | 59.47 ± 11.69 | **130.61 ± 25.24\*\*** | 13.05 ± 6.71 | **0.189 ± 0.56\*** |
| dMSNs + DA | 60.61 ± 11.63 | **122.71 ± 24.41\*\*** | 12.97 ± 5.92 | **0.244 ± 0.16\*** |
| iMSNs | 54.87 ± 13.16 | 117.74 ± 23.47 | 12.55 ± 5.46 | 0.143 ± 0.66 |
| iMSNs + DA | 56.62 ± 14.12 | 118.79 ± 22.55 | 13.72 ± 5.06 | 0.155 ± 0.54 |
| **BIMODAL STIMULATION** | Onset delay (ms) | Peak delay (ms) | Amplitude (mV) | Slope (mV/ms) |
| dMSNs | 23.35 ± 4.83 | 104.97 ± 22.40 | **12.87 ± 6.04\*\*** | 0.400 ± 0.192 |
| dMSNs + DA | 23.46 ± 4.51 | 106.46 ± 24.19 | **16.02 ± 5.16\*\*** | 0.410 ± 0.272 |
| iMSNs | 22.21 ± 6.31 | 97.55 ± 23.06 | 14.04 ± 7.02 | 0.343 ± 0.239 |
| iMSNs + DA | 21.80 ± 5.97 | 97.95 ± 22.91 | 14.13 ± 5.57 | 0.313 ± 0.166 |

To understand how MSNs sensory responses are modulated during the SWO cycle, responses were separated to those occurring during the Up or Down states. In agreement with previous reports, sensory responses of MSNs were robustly induced during the Down states, whereas in the Up states they were characterized by small amplitudes and a high probability of failure (Fig. Supp. 5A-C)^44,50,51^. In fact, there was a significant negative linear relationship between the response amplitude and the MSNs membrane potential (Fig. Supp. 5C, D). Therefore, we focused on the sensory responses evoked during the Down states.

Contralateral whisker stimulation induced responses with an average onset delay of 22.39 ± 3.72 ms and an average peak delay of 95.96 ± 26.14 ms. The average amplitude of these responses was 10.20 ± 5.72 mV, and their slope was 0.163 ± 0.126 mV/ms (Table 2). When comparing the delays, amplitudes and slopes of the whisker responses between dMSNs and iMSNs, no differences were detected. Then, we asked whether DA would modulate tactile responses; nevertheless, neither dMSNs nor iMSNs displayed any significant difference (Table 2; Fig. 3B, C). Thus, we conclude that DA does not have a clear effect on tactile responses.

In comparison, visual stimulation elicited slower responses when compared to tactile ones (mean onset delay difference between single tactile and visual responses: 35.12 ± 12.34 ms, p<0.001; Table 2). Moreover, visual responses displayed an average onset delay of 57.09 ± 12.47 ms, amplitude of 12.76 ± 6.12 mV, peak delay of 123.80 ± 24.82 ms and slope of 0.167 ± 0.085 mV/ms, with no differences between dMSNs and iMSNs in control conditions (Table 2). However, when DA was released, the peak delay of the visual responses on dMSNs decreased (control = 130.61 ± 25.24 ms, DA = 122.71 ± 24.41 ms; p=0.0046; raw data in Table 2, normalized data in Fig. 3D), and this was also accompanied by faster slopes (control = 0.189 ± 0.56 mV/ms, DA = 0.244 ± 0.16 mV/ms, p=0.04). On the other hand, iMSNs did not display any variation when DA was released (Table 2, Fig. 3D). In addition, evoked tactile and visual cortical responses recorded by LFP were analysed and no changes were observed when DA was released (Table Supp. 1, n=12), discarding any effect of DA on S1 and V1.

Finally, to study how DMS-MSNs integrate bimodal information, simultaneous tactile and visual stimuli were delivered. In this evoked synaptic response, the synaptic summation of tactile and visual responses is impeded due to the longer latencies exhibited by visual stimuli compared to tactile ones^44^ (Table 2). Because DA modified the peak delay of visual responses on dMSNs (Fig. 3D), we questioned whether DA could modify the bimodal integration. The analysis of bimodal responses displayed an average onset delay of 22.76 ± 5.58 ms (dMSNs = 23.35 ± 4.83 ms; iMSNs = 22.21 ± 6.31 ms), similar to the value exhibited by single tactile responses (Table 2). This reflects that the early component of the bimodal responses is induced by the whisker input, which presents the fastest onset delays. The population amplitude of the bimodal responses in control conditions was 13.40 ± 6.53 mV, with no differences between dMSNs and iMSNs pathways (Table 2). However, the response amplitude in dMSNs increased ∼25%, from 12.87 ± 6.04 mV to 16.02 ± 5.16 mV (p=0.003) when DA was released, whereas iMSNs were not affected (Table 2, Fig. 3B, E).

In summary, we observed that DA increases the amplitude in dMSNs bimodal responses only; suggesting that this change is facilitated by the peak delay reduction of visual responses, which increases the overlapping time between tactile and visual responses, enhancing the synaptic summation between sensory modalities and promoting their synchronization.

*Ex vivo* reports showed a voltage dependent action of DA on dMSNs ^33,62,65^, increasing their neuronal excitability at depolarized membrane potential. Therefore, DA could affect the amplitude of the dMSNs sensory responses through the activation of D1DRs, when neurons are depolarized. Our visual responses were around 4 mV bigger than the tactile ones and no changes on amplitude were detected after DA release, either on visual or tactile responses (Table 2). However, the delay to the peak of visual responses was reduced. In order to test whether DA is affecting sensory responses directly, we obtained the peak delay differences between control and DA visual and tactile responses and compared them with their respective control amplitudes for dMSNs (Fig. Supp. 6). If the increase of bimodal response is strongly dependent of the direct action of DA on dMSNs, a potential correlation between peak delay differences and control amplitudes must occur on visual responses. However, no correlation was found either for visual or tactile responses, suggesting that the direct action of DA on dMSNs is not enough to explain the increase of amplitude observed on the bimodal responses (Fig. 3).

### Type-specific cortical projection to the dorsomedial striatum

The revealed process arises one question: Why does DA affect only visual responses whereas tactile ones remain unaffected?

To identify a possible anatomical substrate that could explain such specificity, we studied S1 and V1 corticostriatal projections. We first injected biotin dextran amine (BDA) in S1 and V1 of 8 C57BL/6J mice. We observed that DMS received axons from both S1 and V1, although with a considerably smaller covered area by axons from S1 (V1=97 ± 11%; S1=56 ± 11%, p=0.024) (Fig. Supp. 7). In contrast, DLS was densely innervated by axons from S1 (99 ± 4%), and basically empty of projections from V1 (0.5 ± 0.3%) (p=0.022).

Corticostriatal projecting neurons are divided into the pyramidal tract (PT) and the intratelencephalic (IT)^17^ subtypes. Because we observed that DMS receives around half of the innervation from S1 axons than the DLS (Fig. Supp. 7), we wondered whether these two types of cortical neurons in V1 and S1 display different projection patterns to the DMS. To explore this hypothesis, we selectively infected IT or PT neurons in S1 or V1 by injecting the cre-dependent virus AAV2-EF1a-DIO-tdTomato in 8 Tlx3-cre and 8 OE25-cre mice, respectively (Fig. 4A). Our results show that both IT and PT neurons send axons to the DMS from V1 (IT V1 axons = 78.5 ± 4%; PT V1= 34.8 ± 12.5%, p=0.028) (Fig. 4B, C). However, this was not the case for S1 corticostriatal projections: IT S1 neurons projected towards the DMS, while PT projections from S1 were barely detectable or almost absent (IT S1= 51.7 ± 21.7%; PT S1= 4.8 ± 2%, p=0.026) (Fig. 4B, C). Moreover, the PT axons ratio covering the DMS was significantly higher for V1 (PT V1 ratio= 41.52 ± 14.16%; PT S1 ratio= 5.73 ± 2,61%, p=0.028, see methods). The near absence of PT S1 projections confirms that there is a cell-type-specific corticostriatal innervation from S1 to the DMS, while V1 targets the DMS with both PT and IT. Therefore, we asked whether this type-specific projection could explain why only visual responses are affected by DA release.

**Figure 4.**
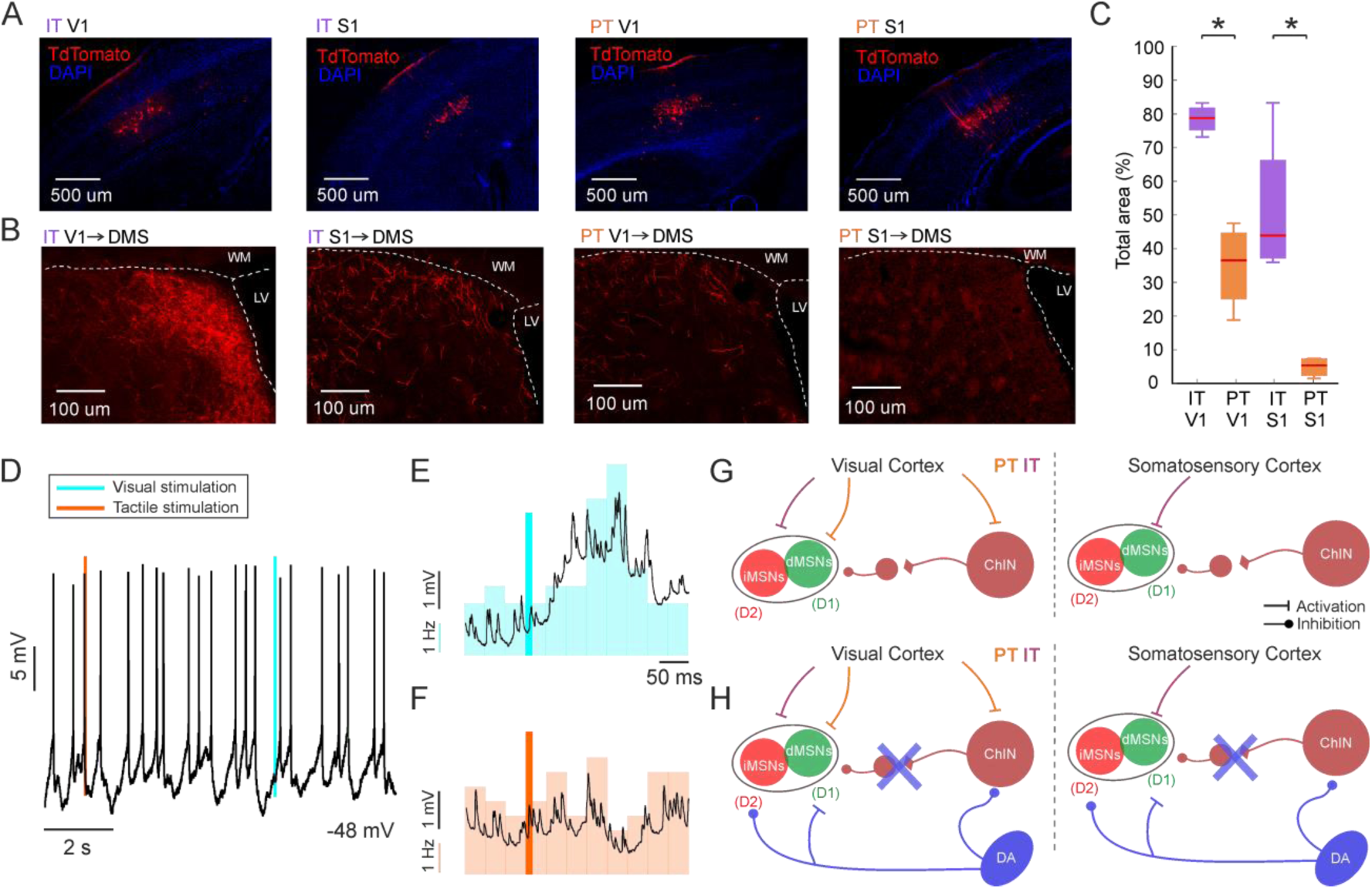
Type-specific corticostriatal projection mediates the synchronization of sensory responses through dopamine release. **A,** Tlx3-cre and OE25-cre mice were injected in layer 5 of S1 and V1 cortices with AAV2-EF1a-DIO-tdTomato-WPRE to target IT and PT neurons respectively. DAPI is shown to verify the cortical layers. **B,** Representative images from each brain shown in A of the axonal coverage of the dorsomedial striatum from IT or PT projections in V1 or S1. **C,** Quantification of the area covered by PT and IT Td-Tomato positive axons from V1 and S1. * p<0.025. **D,** Example of an *in vivo* recorded cholinergic interneuron while performing visual and tactile stimuli. **E, F,** In coloured bars, PSTH of the firing rate after tactile (orange) and visual (blue) stimulation (Bin size = 35 ms). The dark coloured line indicates the sensory stimulation (15 ms). Black line over the PSTH represents the waveform average of the same neuron. **G,** Schematic representation of the experimental set up: *in vitro* whole-cell patch- clamp recordings with viral injections of two different opsins in S1 and V1, while optogenetic stimulation of terminals in the striatum. H, Image showing an example of a recorded cholinergic interneuron, visible thanks to its large soma compared to MSNs. **G, H,** Schematic representation of the corticostriatal interactions in control conditions **(G)** and when DA is released **(H)** in the DMS. Note that disynaptic inhibition from ChINs towards dMSNs and iMSNs is supressed when DA is released.

ChINs are critical regulators of the striatal microcircuits activity due to their large axonal fields^34,66^. In the DMS, ChINs receive massive dopaminergic innervation that inhibits their activity. A recent study showed that, in the DLS, IT neurons fail to recruit ChINs efficiently, when compared to PT neurons^67^. If this is conserved in the DMS, our anatomical result implies that whisker stimulation should have little or no impact onto DMS-ChINs activity due to poor S1 PT to DMS innervation. In addition, ChINs depolarize after whisker stimulation in the DLS ^44^, and they comprise only 1-2% of the total striatal neurons, thus, the probability of recording them during blind *in vivo* patch-clamp recordings in a deep brain region is very low. We recorded 3 ChINs in the DMS that could be clearly distinguished by their electrophysiological properties (Fig. Supp. 1C). In this set of ChINs, we tested the single visual and whisker responses as previously shown (Fig. 4D). In all cases we observed that visual but not tactile stimulation increased ChINs firing rate (Fig. 4E, F). This result is consistent with our anatomical data, as tactile responses fail to recruit ChINs likely due to the lack of PT axons from S1 (Fig. 4B, C), as previously described *ex vivo*^67^. On the other hand, the observed increase in the firing rate of ChINs upon visual stimulation (Fig. 4E), may be explained by the abundant V1 PT to DMS axons that are likely to contact and recruit the DMS-ChINs (Fig. 4C).

Taking all this information together, we propose a model to illustrate how visual inputs can be specifically modulated by DA release (Fig. 4G, H).

Our hypothesis suggests that during resting state, PT V1 axons activate both MSNs and ChINs, while the absence of PT S1 projections makes unlikely for the tactile inputs to recruit ChINs efficiently in the DMS. Therefore, ChINs will increase the inhibition towards MSNs in response to visual but not tactile stimulation ^23,24^ (Fig. 4G). However, when DA is released it will inhibit ChINs^34,42,43,68^ resulting mainly in the disinhibition of MSNs visual responses (Fig. 4H). Due to this disinhibition, visual responses will be “unbraked”, accelerating their responses (Fig. 3D, Table 2). Hence, the peak delay between visual and tactile responses decreases (from 41.33 ± 36.82 ms to 32.09 ± 36.09 ms, p=0.0012, Fig. 3D, Table 2); facilitating the synaptic summation of both inputs in the bimodal stimulation (Fig. 3E). Nevertheless, the effect of ChINs activity blockage towards MSNs does not explain why, after DA release, sensory responses were affected on dMSNs and not on iMSNs. Thus, in order to decipher the proposed functional diagram, we built and tested our hypothesis in a computational model, including the action of DA on both types of MSNs.

### *In silico* dopaminergic control of sensory synchronization

Our model consisted on a population of MSNs, divided in dMSNs and iMSNs cell types, together with a population of ChINs, following the computational models of Humphries et al. 2009^69^ and Gregory Ashby and Crossley 2011^70^, respectively. We tuned the model to fit the evoked firing rates of the ChINs we had recorded *in vivo* (Fig. Supp. 1C). It includes changes in the cell properties of both MSNs types in the presence of DA, as well as the known blockage of ChINs spiking activity when DA is released^42,43,68^.

To characterise the effect of the recruitment of ChINs on the MSNs peak delay, we used the time of their first spike. We stimulated the model using a series of spike trains following a Poisson distribution. We simulated two parallel circuits, both of them consisting in a population of MSNs connected to ChINs and receiving the same input (Fig. 5A). As previously mentioned, IT inputs to ChINs are weaker than inputs from PT neurons^67^. Therefore, based on our anatomical results (Fig. 4), these two circuits differed in the recruitment of ChINs: in one case ChINs were consistently recruited during the stimulation, emulating visual inputs towards the DMS (ChINs*^conn^*), whereas in the other case, ChINs were not triggered, modelling the response to the tactile stimulation (ChINs*^disc^*) (Fig. 5A).

**Figure 5.**
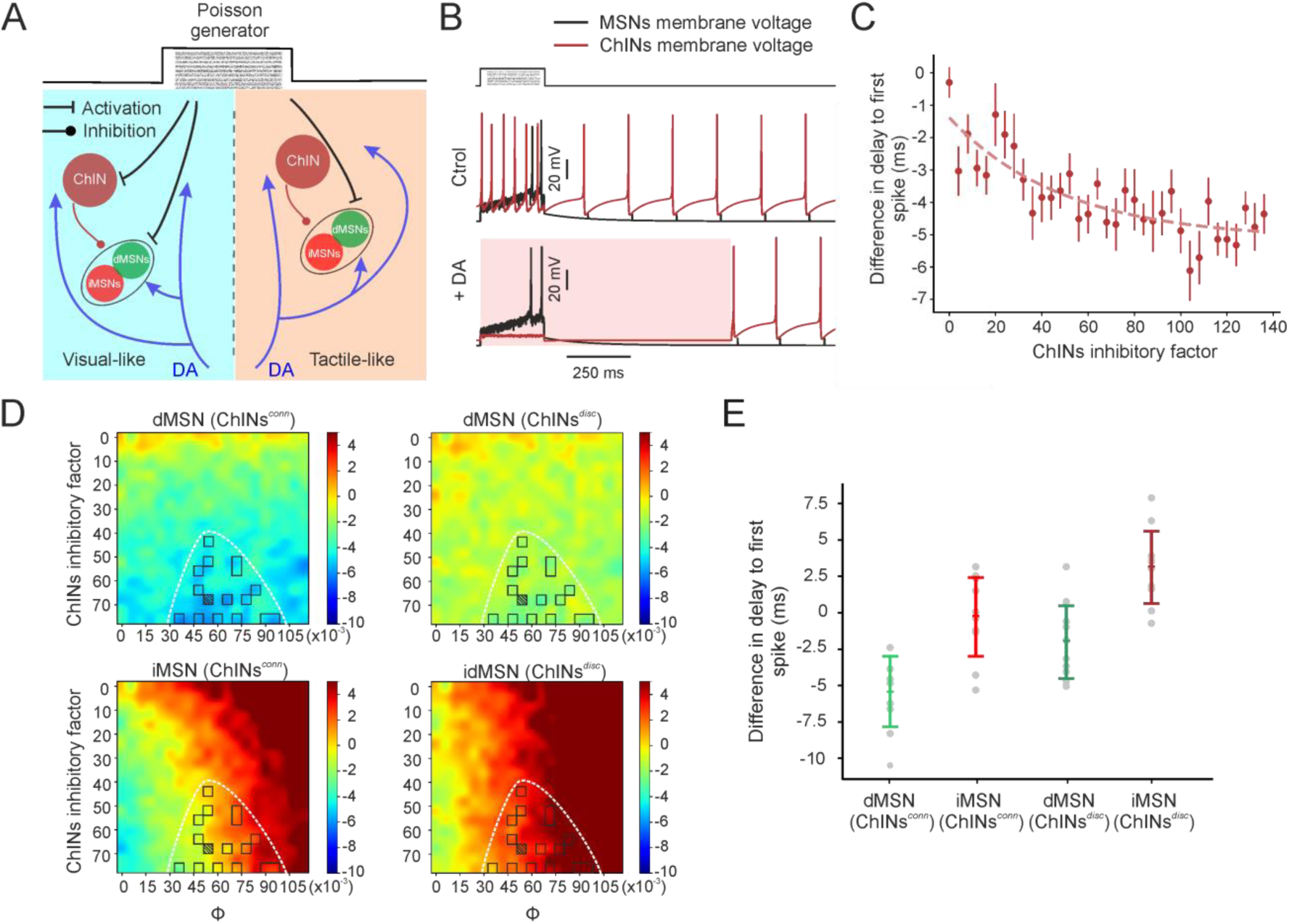
Computational model of corticostriatal connectivity. **A,** Scheme of the computational model. Note how the left population mimics the visual input, while the right population mimics the somatosensory input. **B,** Example traces of evoked responses in a modelled MSN and ChI. Top image corresponds to the basal condition, with no release of DA. Bottom image corresponds to the condition when DA is released (light pink), causing a blocking of the ChIN. **C,** Effects of the ChINs inhibitory factor in the difference in the delay to the first spike of MSNs. An exponential regression (R^2^=0.67) illustrates the dependence between the driving force of the ChIN-MSN synapse and the difference in delay to the first spike of MSNs. Notice how the tendency remains stable once reached a ChINs inhibitory factor of ∼80. **D,** Heat map showing the difference of the delay to the first spike (DA release - basal conditions) in the parameter space that results from the combination of the ChINs inhibitory factor in the range from 0 to 76 and φ (DA modulation) in the range from 0 to 0.114. White shaded line highlights the space in which the delay values were equivalent to the *in vivo* recorded values. Squares correspond to the combination of parameters that produced differences in the delay comparable to the *in vivo* results (mean+/- 1 std of the recorded values). **E.** Representative example illustrating the difference in delay to the first spike (DA release – basal conditions) for dMSNs and iMSNs when ChINs are recruited (ChINs*^conn^*) or not (ChINs*^disc^*) using the values from the shaded square in Fig. 5D (ChINs inhibitory factor = 68; φ = 0.048).

In order to model ChINs inhibition, we used the time constant and delay described here^71^, and explored the driving force of the inhibition of ChINs towards MSNs as a multiplicative free parameter (named “ChINs inhibitory factor”), starting from the strength of a MSN collateral^69^. In each simulation, 20 spike trains coming from the Poisson generator and with a length of 250 ms were delivered to the population. DA release was presented in 50% of the trials with a random distribution, blocking the ChINs activity (Fig. 5B). The delay to the MSNs first spike was compared between trials on which ChINs were or not recruited by the stimulation, and explored the delay difference as a result of the changes in the force of the ChINs-MSN inhibition (Fig 5C). In order to simulate the impact of DA on MSNs, we added the effects onto dMSNs and iMSNs as well as cortico-MSN synaptic currents described here^69^, with exception of the increase of the IKir currents of dMSNs, as we could not observe any change in the membrane voltage during the Down states when releasing DA (Fig. 2F). We explored the space of two free parameters: the driving force of ChINs onto MSNs (ChINs inhibitory factor), and the modulatory strength of DA (“φ”). We tested which combinations of these parameters caused a delay to the first spike that fitted in the mean ± 1 std range of the values of our *in vivo* results for the effect on the peak delay (Fig. 5D). In order to regulate the strength of the dopaminergic modulation of MSNs and following the original model, we tuned the parameter φ, which adjusts the number of open channels and had an original value of 0.8^69^. Even though we do not assume a linear relationship between the effect of DA in the MSNs and the number of open channels, this parameter can still be used to adjust the strength of DA effect to fit the values recorded *in vivo* (Table 2). We found that the parameters space comprised between ChINs inhibitory factor of 40 and 80 (corresponding to 40-80 collateral spikes) and φ values of 0.03 and 0.1 (Fig. 5D, E) produced a difference in the delay to the first spike that was simultaneously consistent with the 4 different stimuli (visual = ChIN*^conn^* and tactile= ChIN*^disc^*; with and without DA release) that were evoked *in vivo*. It is important to notice that the blockage of ChINs is always necessary to reproduce the *in vivo* results, supporting the relevance of this interneuron in the acceleration of the responses of dMSNs, as theorised. Furthermore, the obtained *in vivo* results combined with our computational model explains why the acceleration of the response occurs only in dMSNs, since the disinhibition produced by the blockage of the ChINs is counterbalanced in the iMSNs by their reduction in excitability following DA release.

In summary, we have shown how dopaminergic disinhibition of dMSNs following the blockage of ChINs, together with the activation of D1DRs is a plausible mechanism for the acceleration of the dMSNs peak delay in visual responses, as well as the absence of change in the response to somatosensory inputs.

Our model also provides the parameter space that leads to changes in the delay consistent with our *in vivo* results, depending on the strength of the dopaminergic modulation of MSNs and the driving force of the polysynaptic inhibition of ChINs over MSNs (φ and ChINs inhibitory factor, respectively).

## DISCUSSION

### Dopamine facilitates bimodal sensory synchronization in the dMSNs of the DMS

In the dorsal striatum, as well as in their respective cortical areas, visual responses present longer latencies compared to tactile ones^44^ . This causes a desynchronization between visual and tactile inputs upon simultaneous delivery of these stimuli. A previous study showed that when visual and tactile responses were artificially forced to converge temporally, by delaying the tactile stimulation, the bimodal response amplitude increased^44^. Here, we have observed that this process occurs as a physiological mechanism after DA release.

We found that S1 sends IT but no PT axons to the DMS, while V1 innervates it with both subtypes of cortical projections (Fig. 4). Tracing results also showed that IT V1 corticostriatal axons covered a larger area of the DMS compared to PT V1 axons (Fig. 4C), supporting the idea that PT neurons are smaller in number and have less complex axonal tree than IT neurons^18,72,73^. In the DLS, PT neurons recruit ChINs more efficiently than IT neurons, due to a stronger relative input towards ChINs^67^. Accordingly, in few examples during *in vivo* recordings we observed that ChINs increase their firing when responding to visual, but not to tactile stimulation (Fig. 4E, F), suggesting that the IT/PT connectivity imbalance to ChINs is also present in the DMS.

DA release correlates with the pauses of ChINs activity, as it has been extensively described^42,43,68,74^. In the DMS, DA pauses ChINs firing during approximately one second^42^. This large pause results in the disinhibition of dMSNs. In addition, recent results report that ChINs synchronize their activity, suggesting a network action over the striatal microcircuits^25^.

An important point arising from our results refers to why this DA-dependent synchronization is detected exclusively in dMSNs but not iMSNs. Both types of MSNs are also affected by the activation of their respective dopamine receptors. *Ex vivo* experiments showed that DA regulates their excitability: the activation of D2DRs expressed in iMSNs results in reduced excitability and responsiveness^35,37,42,68^. This is supported by the output of our *in silico* model, which indicates that the reduction in the excitability of iMSNs mediated by DA can counterbalance the disinhibition caused by the blockage of ChINs, resulting in the absence of changes on visual responses after DA release, in agreement with our biological results (Fig. 5D, E). On the other hand, DA increases the excitability of dMSNs without affecting their input resistance, even when synaptic transmission and neuromodulation is blocked. This suggests a direct action of DA on dMSNs by the activation of D1DRs receptors^37^. D1DRs change cellular function via adenyl cyclase (AC) and protein kinase A (PKA) activation by increasing Cav1 L-type Ca^+2^ channel currents and decreasing somatic K^+^ A_s_-currents^33,75^, thus promoting Up state transitions and excitatory synaptic potentials^36,75,76^. Finally, under similar conditions to our *in vivo* experiments it was shown that DA release increases PKA levels in dMSNs, while no changes were detected in iMSNs of the nucleus accumbens^77^. In summary, the compelling evidences suggest that DA promotes the activation of dMSNs, making them more predisposed to respond to sensory stimuli.

The last prediction of our *in silico* model suggests that a very small release of DA can support the reduction of the delay of the response in both dMSNs and iMSNs (Fig. 5D), as long as DA is still able to block ChINs. This is due to the impact that the concentration of DA has on the delay to the first spike of iMSNs, combined with the stable effect on dMSNs delay, which is mostly modulated by (absence of) ChINs’ recruitment. Larger increases of DA concentration will hamper iMSNs, with little effect on dMSNs. Interestingly, *ex vivo* experiments have shown that an increase of DA concentration from 60 µM to 120 µM does not result in a higher excitability of dMSNs, while it decreases the excitability of iMSNs^37^. In our model, this predicts that the relative timing of dMSNs and iMSNs, in response to a given input, can be modulated by the amount of DA released in the DMS, hence providing a mechanism to regulate the timing of the output of these pathways. In this sense, it has been reported that the co-activation of both MSNs pathways is important for action selection and initiation^78^. Therefore, this mechanism may need the normal responsiveness of iMSNs, for instance to inhibit competing motor programs.

Other striatal interneurons could be involved directly targeting MSNs after sensory stimulation; such as fast spiking interneurons (FSIs), which respond to whisker and visual stimuli in the DMS^44^ and innervate both types of MSNs with high probability of connection^79,80^. FSIs increase their activity by DA, mediated by the D5DRs receptor activation^81^. However, FSI completely avoid ChINs^82^. On the other hand, low threshold spike interneurons (LTS) receive monosynaptic excitatory inputs from cortex^83,84^. Because DA depolarizes LTS^85^ and they send feedforward inhibition to MSNs, its output would inhibit MSNs. In addition, a recent study showed that the axonal projections from pyramidal neurons of S1 to striatal LTS is sparse or absent, being copious from M1^86^. Lastly, TH interneurons (THINs) also receive monosynaptic glutamatergic inputs from cortex^75^. Bath application of DA to THINs results in an enhancement of their excitability, which generates an increase of direct inhibition to MSNs^75^. In summary, with exception of the ChINs, evidences suggest that DA increases the excitability of different types of the striatal interneurons. However, we did not find any significant decrease of amplitude and slope or increase on delays of the tactile and visual responses after DA release (Table 2), suggesting that they are not playing a pivotal role on the modulation that DA has on our sensory responses in the DMS.

It can be possible that the presynaptic inhibition between MSN-MSN connections would result in the disinhibition of postsynaptic dMSNs. Nonetheless, the probability of connection between MSNs is sparse and characterized by a low and variable release probability^79,87,88^; and both MSNs receive excitatory synaptic inputs from PT and IT pyramidal cells^89^. In addition, our results showed that both types responded to visual and whisker stimulation similarly in control conditions (Table 2). Therefore, if the disinhibition observed on dMSNs after DA release was mediated by a MSN-MSNs connection, this would likely occur to both visual and whisker responses. However, changes were only detected for visual responses.

In any case, we cannot rule out this or other possibilities and future studies are necessary to increase the detail level of our model.

### DA release decreases high frequency oscillatory energy in MSNs during Up states

In the BG, exaggerated beta-band oscillations power has been established as one of the signatures of Parkinson disease (PD)^90–94^, and its appearance is usually taken as a sign of DA depletion^90,95^. On the other hand, this exaggerated beta-band oscillatory activity is reduced by L-DOPA treatment in PD patients^90,92–94,96^. During cortical Up states, MSNs, ChINs and other striatal interneurons receive barrage of synaptic inputs that induce high frequency oscillations^15,97,98^. In addition to glutamatergic transmission from cortex and thalamus, ChINs also promote glutamate release to MSNs, generating a delayed second phase of excitation^67^. This could induce the high frequency oscillatory activity detected during the Up states on MSNs. Therefore, the blockage of the ChINs activity by DA release could mediate the decrease of this high frequency oscillatory energy detected in both types of MSNs during Up states (Fig. 2B). Moreover, as discussed, the excitability increase that DA induces on diverse types of interneurons could also mediate this reduction.

Finally, this high frequency reduction could also be mediated by GABA co-release generated by activation of dopaminergic neurons^36,59–64^. A couple of recent *ex vivo* studies showed that no glutamate co-transmission was observed in MSNs in the DMS^99,100^; whereas a weak co-release of GABA was documented^99^. However, in our hands we did not identify any fast hyperpolarization or depolarization in MSNs after the optogenetic or electrical stimulation of DA neurons (Fig. Supp. 4A, B). *Ex vivo* studies often reported stronger effects on short-term plasticity than *in vivo*, due to the larger probability of release described *ex vivo*^101^, suggesting discrepancies on synaptic transmission between preparations. This can be mediated by differences in the ionic environment, with lower concentration of Ca^2+^ and Mg^2+^ and higher K^+^ *in vivo*^102^. In addition, during *ex vivo* recordings synaptic activity is highly decreased and the release probability depends in a complex way on recent activity. For instance, the emergence of SWO reduces the levels of short-term depression^103^. Other factors affecting synaptic transmission are temperature^104^ (often room temperature in vitro vs. physiological *in vivo*), or maturational differences^105^ (young vs. adults). All these factors could induce discrepancies on synaptic transmission, affecting the levels of GABA co-release, being probably very low during the SWO and undetectable in our hands.

### Functional significance

Previous reports demonstrated that the combination of multimodal stimuli facilitates an adequate response to relevant events^106–111^. However, still it is unclear how this occurs, since the different latencies when processing visual and tactile inputs result in a delay which causes the independent processing of these sensory inputs, as observed in DMS-MSNs as well^44^. A similar temporal asynchrony between sensory modalities has been described in motor control studies, in which visual motor feedback is tens of milliseconds slower than the tactile one^112,113^. How the brain operates with these discrepancies of sensory latencies is an essential question in order to understand how visual and somatosensory perturbations engage motor responses^114^. In addition, the temporal processing of sensory data is abnormal in Parkinsonian patients^115^, and impaired detection of paired multisensory information of PD patients has been documented^9^. Moreover, these deficits improve under dopaminergic drugs^9,116,117^. In this work, we found that DA enhances the sensory synchronization by controlling the timing of MSNs responses and proposed a new mechanism underlying this process mediated by the release of DA. This can contribute to the understanding of sensory-motor interactions that occur during fine movements and provide an explanation of how DA can be related with some sensory deficits.

## ACKNOWLEDGEMENTS

We thank Dr. Gilad Silberberg (Karolinska Institutet) for generous donation of D2-Cre(ChR2)- YFP mouse and for helping us to start-up the laboratory. We thank Dr. Rui M. Costa for his generous contribution with transgenic mice, viral vectors, and lab facilities. We thank the Dr. Tian laboratory for their generosity sharing the dLight1.2-EGFP expressing vector for our initial experiments. The study was supported by MINECO Starting Grant I+D Jovenes Investigadores [BFU2014-60809-IN], the CSIC-Severo Ochoa Excellence Programmes of the Instituto de Neurociencias [SEV-2013-0317 and SEV-2017-0723] and the CSIC Interdisciplinary Thematic Platform (PTI+) NEURO-AGING+ (PTI-NEURO-AGING+) and the [MICINN PID2021-129070NB-I00]. M.S. was supported by the MINECO Fellowship [BES-2015-072187] and by the PTI- NEURO-AGING+. J.A-C is currently supported by a Margarita Salas Fellowship [2021/PER/00020] funded by Ministerio de Universidades, UMH and European Union. N.A.M. was supported by Fundação para a Ciência e a Tecnologia [SFRH/BPD/88309/2012]. R.d.l.T-M was supported by a Karolinska Institute postdoctoral scholarship.

## AUTHOR CONTRIBUTION

M.S. performed the electrophysiological and fiber photometry experiments, viral injections and BDA tracing, formal analysis, and the writing, review and editing of the original draft. J.A-C. performed formal analysis, built the computational model and carried out the writing, review and editing of the original draft. N.A.M. performed viral tracing and review and editing of the original draft. C.G-F. provided methodological assistance and performed the review and editing of the original draft. R.d.l.T-M. performed electrophysiological experiments and the review and editing of the original draft. R.R. conceptualized, provided resources and funding, supervised and performed the writing, review and editing of the original draft.

## METHODS

### I. Animals

#### Ethical approval

All the experimental procedures were conformed to the directive 2010/63/EU of the European Parliament and of the Council, and the RD 53/2013 Spanish regulation on the protection of animals use for scientific purposes, approved by the government of the Autonomus Community of Valencia, under the supervision of the Consejo Superior de Investigaciones Científicas and the Miguel Hernandez University Committee for Animal use in Laboratory.

#### Animal model

The total amount of mice was 52 animals, of both sexes and between 2 and 8 months of age. D2-cre x ChR2 x DAT-cre mice were used to distinguish dMSNs and iMSNs during our recordings, as previously described^15,51^ (Fig. 1E). Briefly, the expression of ChR2 specifically in iMSNs allowed their identification by optogenetic stimulation. At the same time, the expression of cre in DAT neurons of the same animals was used to selectively express ChrimsonR in dopaminergic neurons of SNc^118^ to stimulate the release of DA (n=15 animals, 12 + 3 control) (Fig. 1C). To obtain this mouse line, D2-Cre (ER44 line, GENSAT) mice were crossed with the Channelrhodopsin (ChR2)-YFP reporter mouse line (Ai32, the Jackson laboratory) to induce the expression of ChR2 in iMSNs. The resulting mouse line was then crossed with DAT-cre mouse (the Jackson laboratory). The expression of ChR2 and ChrimsonR was checked in all the brains and no DAT-ChR2 axons were found (Fig. 1C). Moreover, no functional interaction occurs between the 465 nm and the 660 nm led activation; as no changes in the SWO properties were observed after stimulation with 660 nm. On the other hand, the 465 nm LED stimulation was only used for the MSNs identification, at the beginning of every recording during few seconds (more information above, Methods V section).

Additional animals from the DAT-cre mouse line were used to perform fiber photometry experiments (n=13 animals (7 + 6 control, Fig. Supp 3)). C57BL/6J mice (n= 8 animals) were used for BDA injections (Fig. Supp. 7A). OE25-cre (Tg(Chrna2-cre), OE25Gsat/Mmucd, GENSAT) (n=8 animals) and Tlx3-cre (Tg(Tlx3-cre)PL58Gsat/Mmucd, GENSAT) (n=8 animals) mice lines were used to perform PT and IT selective axonal tracing respectively from S1 and V1 cortices to the dorsal striatum (Fig. 4).

### II. Viral tracing

#### Viral targeting in SNc

In order to induce DA release by optogenetic stimulation, we infected the dopaminergic neurons in the SNc. Isofluorane anesthetized D2-cre x ChR2 x DAT-cre mice were immobilized in a Stereotaxic Alignment System (Kopf Instruments). AAV5-hSyn-FLEX-ChrimsonR-tdTomato (n=12 animals) or AAV5-CAG-FLEX-tdTomato (n=3 animals) (both from UNC Vector Core with titers of at least 1x10^12^ vg/ml), were uni-laterally injected intracerebrally (500 nL) with an injector (Nanoliter, WPI) into the SNc using the following coordinates: AP -3 mm, LM 1.5 mm, DV -3.5 mm^119^. Experiments were performed at least 8 weeks after the viral injection.

#### Viral targeting in DMS

In order to measure DA release in the DMS using fiber photometry, DAT-cre mice (n=7 animals + 6 control animals) were first injected in the SNc, as described above. Then, 6 weeks later, a second injection was performed unilaterally in the DMS, using the following stereotaxic coordinates^119^: AP 0 mm, LM 2 mm and 3 depths below the surface (DV -1.6 mm, -1.9 mm, and -2.2 mm; 100 nl at each depth, total 300 nl), with AAV5-hSyn-dLight1.2-EGFP, obtained from Dr. Lin Tian laboratory (University of California Davis, USA) and Addgene. Experiments were performed at least 2 weeks after the viral injection.

#### Biotin Dextran Amine tracing in S1 and V1

In order to trace the projections from S1 and V1 towards the striatum, C57BL/6J mice (n=8) were anesthetized and immobilized as described above. Biotin Dextran Amine (BDA) (Sigma Aldrich) was injected unilaterally in S1 (following coordinates: AP -1.5 mm, LM 3.25 and 3.75 mm, DV -0.8 mm), and V1 (following coordinates: AP -3.75 mm, LM 2.5 and 2.75 mm, DV -0.8 mm)^119^, injecting 150 nl in each lateromedial coordinate to cover the largest area as possible. For the anatomical study, mice were sacrificed 10 days after the injection by receiving an overdose of sodium pentobarbital (200 mg/kg I.P.).

#### Viral tracing in S1 and V1

In order to selectively target PT and IT cortical neurons projecting towards the striatum, Tlx3- cre (n=8) and OE25-cre (n=8) mice were anesthetized and immobilized as described above and 300 nl of AAV2.EF1a.DIO.tdTomato.WPRE virus (UNC Vector Core with a titre of at least 1x10^12^ vg/ml) were injected unilaterally in S1 and V1 following the same coordinates as described above for AAV tracing to cover the largest area as possible. Sacrifice was performed 31 days post injection by receiving an overdose of sodium pentobarbital (200 mg/kg I.P.).

### III. Electrophysiology

#### *In vivo* Electrophysiological recordings

D2-cre x ChR2 x DAT-cre mice of both sexes (n=15, n=9 males and 6 females), were used to perform the whole-cell recording experiments (Table Supp. 2). Anaesthesia was induced by intraperitoneal injection of ketamine (75 mg/kg) and medetomidine (1 mg/kg) diluted in 0.9 % NaCl. A maintaining dose of ketamine (30 mg/kg i.m.) was administrated every 2 hours or after changes in the EEG or reflex responds to paw pinches. Tracheotomy was performed to increase mechanical stability during recordings by decreasing breathing related movements. Mice were placed in a stereotaxic device (customized Stoelting stereotaxic base) and air enriched with oxygen was delivered through a thin tube placed 1 cm from the tracheal cannula. Temperature was maintained at 36.5 ± 0.5°C using a feedback-controlled heating pad (FHC Inc.). Craniotomies were drilled (S210, Camo) at several sites from bregma: AP 0 mm, LM 2.5 mm (DMS); AP −1.5 mm, LM 3.25 mm (S1); AP −3.5 mm, LM 2.5 mm (V1)^119^. Animals were sacrificed after the experimental session by receiving an overdose of sodium pentobarbital (200 mg/kg I.P.).

#### *In vivo* whole-cell recordings

Whole-cell patch-clamp recordings were obtained from the DMS between 2050 and 2567 µm deep in a perpendicular penetration angle of ∼30°. The exposed brain was continuously covered by 0.9% NaCl to prevent drying. Signals were amplified using a MultiClamp 700B amplifier (Molecular Devices) and digitized at 20 KHz with a CED acquisition board and Spike 2 software (Cambridge Electronic Design). Borosilicate patch pipettes (1B150F-4, WPI), were pulled with a Flaming/Brown micropipette puller P-1000 (Sutter Instruments) and had an initial resistance of 6-12 MΩ, with longer tips than the standard ones to minimize cortical damage. Pipettes were back-filled with intracellular solution containing: 125 mM K-gluconate, 10 mM KCl, 10 mM Na-Phosphocreatine, 10 mM HEPES, 4 mM ATP-Mg and 0.3 mM GTP-Na. pH and osmolarity were adjusted to ∼7.4 and ∼280 mOsm/L, respectively. Biocitin (0.2-0.4%, Sigma Aldrich) was then added to the intracellular solution to reconstruct the recorded cell after every experiment (Fig. 1B). For the analysis of the spontaneous activity, 100 s of spontaneous activity (no current injection, no stimulation) were used from the recording. To analyse the dopaminergic modulation of spontaneous activity, 100 s of the recording without stimulation were compared to 100 s of the same recording in which DA was optogenetically released every 5 s to avoid brain state entrainment. 420 s of the recording with optogenetic and sensory stimulation were used to analyse the DA modulation on sensory responses, ensuring for each recording a minimum of 14 events for each condition/cell. For all the neurons, input resistance (Table 1) was measured as the slope of a linear fit between the injected current steps and membrane potential of 2 s duration. In order to quantify the described inward membrane rectification of MSNs^120,121^, mean resistance during Up or Down states was analysed in response to negative and positive current steps (Table 1). Neurons with resistances < 100 MΩ were excluded from the analysis. The time constant (tau) was calculated as the time required for the membrane voltage change to reach the 63% of its maximum value. Capacitance was obtained by dividing the time constant between the voltage membrane resistance. Neurons with a resting membrane potential above -50 mV and/or having a deviation by more than 10 mV from their initial resting membrane potential were excluded from the analysis. The action potential threshold for our cells was determined to be -40 mV, therefore, the firing rate was calculated by using a threshold in the membrane potential of the cell (-20 mV) and counting the number of action potentials presented in a given time at that membrane potential. The SWO frequency was calculated as the number of Up states in a given recorded time. From the 38 recorded neurons, 33 were identified as MSNs by their electrophysiological properties and morphology (Fig. 1B, D). From the remaining 5, 2 were identified as FSIs and 3 as ChINs by their electrophysiological properties. FSIs tend to have higher input resistance and an increased firing rate compared to MSNs; while ChINs display a tonic firing and voltage sag response (Fig. 1G-I, Table 1, Fig. Supp. 1). The average recording time for all MSNs was 49.96 ± 10.27 min (minimum=23 min, maximum=72 min; n=33)). As the average recording time was quite variable, not in all neurons was possible to accomplish all the experimental protocols. First, recorded MSNs were optogenetically identified and classified as a putative dMSN or iMSN by the means of the optopatcher^122^ (Fig. 1E); second, we studied their electrophysiological properties by injecting positive and negative intracellular current steps. Afterwards, we aimed to study the high frequency oscillatory activity during their spontaneous activity and the impact of DA on visual and tactile responses. Hence, from the 33 neurons identified as MSNs, all of them (18 dMSNs and 15 iMSNs) were identified and used to study their electrophysiological properties, 28 (18 dMSNs and 10 iMSNs) were analysed to study the effect of DA onto the high frequency oscillatory activity, and 29 (14 dMSNs and 15 iMSNs) were included to study the effect of DA in sensory processing.

ChINs displayed their characteristic voltage sag response to current step injections (7.78 ± 2.20 mV), depolarized membrane potential (-51 ± 6.55 mV) (Table 1), and spontaneous discharge activity (1.35 ± 0.41 Hz) (Fig. Supp. 1A, B). On the other hand, FS showed a high firing rate (2.69 ± 0.95 Hz) and hyperpolarized membrane potential (-65 ± 7.07 mV) (Table 1).

#### *In vivo* extracellular recordings

Extracellular recordings were obtained using unipolar tungsten electrodes with impedances of 1-2 MΩ to confirm that sensory stimulations elicited responses. The electrodes were placed in infragranular layers (1000 µm depth from the pia) of S1 and V1 with an angle between 15° and 25°. Recordings were amplified using a Differential AC Amplifier model 1700 (A-M Systems) and digitized at 10 KHz with CED and Spike-2 simultaneously with the whole-cell recording. Evoked responses where observed and afterwards quantified.

### IV. Stimulation protocols

In all the cases, the different stimuli were randomized, with an interstimulus interval of 5 s (0.2 Hz). For each cell, tactile, visual and bimodal stimuli were launched during a recording with a minimum duration of 420 s, ensuring at least 14 events for each stimulation/cell. Sensory stimulations were randomly preceded (30 ms) by an optogenetic activation of DMS dopaminergic terminals in the 50% of cases. However, due to the longer duration of the Down states with respect to the Up states, the stimulus occurred during the Down states in most of cases (Fig. Supp. 5A, B).

#### Tactile stimulation

The displacement of the main whiskers was obtained by brief air puffs (15 ms of duration, 20 p.s.i.) by means of a Picospritzer unit (Picospritzer III, Parker Hannifin, NJ), via 1 mm diameter plastic tubes, placed at ∼50 mm in front of the contralateral side of the snout. The latency between the computer command and the whisker movement was measured using an extracellular tungsten electrode placed in one of the central whiskers and was determined to occur 20.05 ± 0.1 ms (n = 5 animals) following the trigger command. Therefore, the reference onset time was determined as 20 ms following the computer trigger command. Tactile responses were confirmed by monitoring the activation of the ipsilateral S1 using LFP recordings.

#### Visual stimulation

A brief visual stimulation (15 ms of duration) was delivered by a white light LED positioned ∼50 mm from the contralateral eye. The eye was covered with artificial eye drops (Viscotears, Bausch+Lomb, Germany) in order to prevent drying, as previously described ^123^. Visual responses were confirmed by monitoring the activation of the ipsilateral V1 using LFP recordings.

#### Bimodal stimulation

Tactile and visual stimuli were delivered simultaneously using the same protocols as described above.

### V. Optogenetics

#### Optogenetic identification of *in vivo* recorded neurons

In order to identify “on line” the specific type of MSNs belonging to the direct and indirect pathways, the optopatcher was used^15,51,122,124^ (A-M systems, WA USA). Pulses (SLA-1000-2, Two-channel universal LED driver, Mightex systems) of blue light (Fiber-coupled LED light source FCS-0470-000, peak excitation wavelength of ∼470 nm, Mightex systems) controlled through Spike 2 software were delivered using an optic fiber (200 µm diameter, handmade) inserted into the patch-pipette, while recording their spontaneous activity (Fig. 1E). One or two serial pulses with 5 light steps of 500 ms each were delivered every 2 s with increasing intensity from 20% to 100 % of full LED power (minimal light intensity 0.166 mW; maximal intensity 0.83 mW at the tip of the fiber). Power light was measured with an energy meter console (PM100D, Thorlabs). Positive cells responded to light pulses by depolarizing their membrane potential. Positive cells (Fig. 1E, green trace) responded within 2.73 ± 1.29 ms of latency (ranging from 0.8 to 5 ms) to light pulses, by a step-like depolarization of 10.22 ± 8.33 mV (ranging from 2.9 to 19.6 mV) at maximal stimulation intensity. Negative cells did not show any depolarization to light pulses (Fig. 1E, black trace).

#### Optogenetic activation of dopaminergic axons in the DMS

Optogenetic activation was done placing an optic fiber of 200 µm in the vicinity of the recording region with a 90° angle connected to a light source (high power LED Prizmatix) (Fig. 1A). Light intensity was 2.4 mW at the tip of the fiber. Based on previous descriptions^125–127^, we designed an optogenetic train with 4 light pulses of 15 ms at 15 Hz (interpause of 66 ms, peak excitation wavelength of ∼660 nm), that mimicked the burst frequencies of action potentials discharge of dopaminergic neurons^125,127,128^ (Fig. Supp. 3C). This optogenetic stimulation was randomly launched every 5 seconds and 30 ms before tactile, visual or bimodal stimulation or without sensory stimuli.

#### Optogenetic-induced dopamine release measurement with fiber photometry

In order to understand the specific impact of DA onto DMS-MSNs activity, we first compared the activation of the dopaminergic terminals in the DMS with the stimulation of cell somas in the SNc. Experiments were carried out in 7 DAT-cre animals in which the SNc was injected with AAV5-hSyn-FLEX-ChrimsonR-tdTomato^118^ and in 6 DAT-cre control animals injected with AAV5- CAG-FLEX-tdTomato. Additionally, six weeks later, all animals were injected with AAV5-hSyn- dLight1.2-EGFP^54^ in the DMS as previously described (Fig. Supp. 3A, B). Fiber photometry experiments were then performed similarly as in previous reports^53,54^. Briefly, for colour imaging, the 1-site 2-colour Fiber Photometry system was used (Doric Lenses): The 465 nm wavelength was used to capture the fluorescence changes related to the dLight1 sensor, whereas the 405 nm wavelength was used as isosbestic reference point to detect artefactual changes not related with the sensor^54^. The fluorescence captured from the 405 nm signal remained constant during our optogenetic stimulations (Fig. Supp. 3C, black traces), ensuring that the fluorescence increase observed in the 465 nm signal was not an artefact (Fig. Supp. 3C, blue traces). Acquired photometry data from the 405 and 465 nm channels were processed with custom codes written in Matlab. Raw data from each channel was down-sampled and low-pass filtered at 25 Hz using a 2nd order Butterworth filter. To calculate the changes in fluorescence to yield the ΔF/F PSTHs, a linear fit was applied to both signals and afterwards we aligned them. Then, the fitted 405 nm signal was subtracted from the 465 nm signal. The resultant signal was then divided by the fitted 405 nm signal to yield the ΔF/F values. Afterwards, the ΔF/F values were smoothed and aligned to the stimulation trigger to compute the PSTHs. To induce the optogenetic release of DA, the stimulation train consisted on 4 red light pulses (peak excitation wavelength of ∼660 nm), each of them with a duration of 15 milliseconds at 15 Hz (Fig. Supp. 3C), mimicking the experimental protocols of previous studies^125,127,128^. The optogenetic stimulation of dopaminergic neurons in the SNc increased the levels of fluorescence related to DA to 2.29 ± 0.10% in the DMS (Fig. Supp. 3D), a value similar to the ones reported previously^54^. Next, we repeated the experiment but stimulating the dopaminergic terminals directly in the DMS. In this case, the fluorescence increased to 0.56 ± 0.25% (Fig. Supp. 3D), approximately a quarter of the amount obtained when stimulating the SNc (p=0.002). In both cases, the onset of the fluorescence responses was similar between stimulations (Onset delay: SNc = 62.4 ± 12.03 ms, striatal terminals = 55.57 ± 10.01 ms, p=0.67, Fig. Supp. 3E). In all cases, viral infection was anatomically confirmed by the expression of the reporters TdTomato and EGFP from the ChrimsonR and dLight constructs, respectively (Fig. Supp. 3B). Finally, we performed a set of control experiments, in which AAV5-CAG-FLEX-tdTomato was injected in the SNc of 6 DAT-cre mice [see methods]. In this case, changes in fluorescence related to DA release were undetectable (0% of fluorescence increase) either when stimulating the axonal terminals or somas (Fig. Supp. 3C, D). These experiments demonstrate that optogenetic stimulation of the SNc or the DMS is a reliable method to release DA in the striatum. However, the stimulation of DA neurons in the SNc results in a release of DA not only in the DMS, but also in other striatal subregions, such as subthalamic nucleus^129^, external and internal globus pallidus^130,131^, as well as to other brain regions. Henceforth, in order to increase the precision of DA release, we used the stimulation of axons in the DMS to study the effects of DA onto MSNs sensory responses and spontaneous activity.

### VI. Histology

#### Morphological reconstruction

At the end of each *in vivo* experiment the mouse was sacrificed with a lethal dose of sodium pentobarbital (150 mg/Kg) and perfused with a solution containing 4% paraformaldehyde in 0.1 M phosphate buffer (PB, pH 7.4). Brains were extracted and stored in PBS solution until the cutting. Before cutting, brains were transferred into PBS containing 30 % sucrose for 24/48 hours. Coronal or sagittal slices (20 µm thick) of both hemispheres containing the entire striatum from the recorded side (from AP 1.4 mm to AP -1.3 mm, following Paxinos & Franklin ^119^ were obtained using an automatic digital criotome (Microm) and collected on gelatin coated slides (ThermoFisher). Sections were incubated over night with Cy3-conjugated streptavidin (Jackson Immuno Research Laboratories) diluted (1:1000) in 1 % BSA, 0.3 % Triton-X 100 in 0.1 molar PBS. Finally, the gelatin coated slides were covered with mowiol (Calbiochem) and mounted with coverslips (ThermoFisher). Recorded and filled neurons were then reconstructed using a fluorescence microscope (DM 6000B, Leica) and a camera (DC350 FX, Leica) (Fig. 1B).

#### BDA, PT and IT axonal tracing

At the end of each experiment, the mouse was sacrificed and perfused, and the brain was stored as mentioned above. Before cutting, brains were transferred into PBS containing 30% sucrose for 24/48 hours. Coronal slices (40 µm thick) of both hemispheres from the entire brain (BDA brains, n=8) (PT and IT brains, n=16), were obtained using an automatic digital criotome and collected on gelatin coated slides as described above. BDA slices were incubated over night with Cy3-conjugated streptavidin diluted (1:1000) in 1 % BSA, 0.3 % Triton-X 100 in 0.1 molar PBS. PT and IT slices were not incubated. Afterwards, all slices were incubated 20 min at RT with DAPI (Sigma). Finally, the glass slides were covered with mowiol (Calbiochem) and mounted with coverslips (ThermoFisher). To perform the anatomical study, we took images in half of the slices containing the DMS and DLS from the recorded side (from bregma: AP 1.4 mm to AP -1.3 mm, following Paxinos & Franklin^119^. Images were obtained with a fluorescence microscope (DM 4000B, Leica) and a camera (Retiga 2000R, Qimaging). Each taken image covered an approximate area of ∼0.13 mm^2^ (approximately 360 µm per side) in the DMS or DLS (Fig. 4B; Fig. Supp. 7A). We also took images of the injection place in S1 and V1 in each of the brains to ensure that the number of infected cells and the fluorescence expression was similar between brains (Fig. 4A). In order to estimate the density of cortical projections, we binarized and applied to each image a threshold computed as the mean fluorescence intensity plus 6 ± std of all the images, using custom Matlab code. We then computed the total area of pixels above the threshold, which corresponded to the area covered by the labelled axons. PT ratio from V1 and S1 was calculated as follows for each cortical region, where PT (DMS area covered by PT axons); IT (DMS area covered by IT axons):

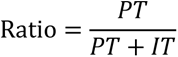

### VII. Computational model

We built our model based on the canonical spiking model of^132,133^ to test our hypothesis regarding the reduction of the peak delay due to a disinhibition of MSNs mediated by a dopaminergic blockage of ChINs.

The model consisted on 200 MSNs (Eq. 1), 100 dMSNs and 100 iMSNs, reproducing the reduced model of ^69^ and 4 ChINs (Eq. 2), implemented as in^70^. MSNs from the direct and indirect pathway only differed in their response to DA. Capacitance of ChINs was adjusted to the values obtained in our recordings *in vivo*. We tuned the constant value in the ChINs voltage derivative from 950 to 346, in order to fit the spontaneous firing rate of ChINs to ∼3 Hz, as described *in vivo*^134,135^. Capacitance (C) of MSNs = 15.2, while C of ChINs = 32. The parameter k has a value of 1 for MSNs and 1.2 for ChINs.

MSNs (Eq. 1)

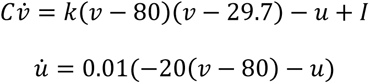

If v≥-30mV then v -55mV, u u+d, with d=91 TANs (Eq. 2)

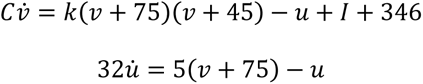

If v≥-30mV then v -56mV, u u+250

#### Circuit connectivity

Stimulation of MSNs and ChINs was modelled following a Poisson process with an event rate of 15 KHz when active. Corticostriatal and ChINs inhibition of MSNs synaptic dynamics were modelled following^69^ (Eq. 3-7) and included AMPA, NMDA and GABA currents (Eq. 3-7).

Synapses (Eq. 3-7)

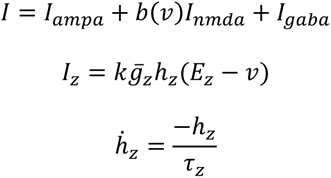

When a presynaptic spike arrives, 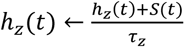

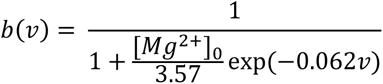

Activation of MSNs was tuned to produce robust depolarization but scarce firing rate (1-2 spikes), and activation of ChINs was tuned to evoke ∼6 spikes in 250 ms, as shown in the traces recorded *in vivo* (Fig. Supp. 1C). This was done adjusting the parameter S in the table shown below and applying a correction factor to multiply the input current to MSNs and TANs that was evoked by the Poisson generator. This correction factor had a value of 0.297 for MSNs and 0.437 for TANs.

In order to model the polysynaptic inhibition of ChINs onto MSNs, we started from the same synaptic dynamics used for MSN-MSN collateral inhibition in^69^. Then, we assigned a value of 15 ms to the synaptic delay and a time constant of 120 ms, following the results of^71^. At last, we tuned the conductance starting from the original conductance of the MSN-MSN synapses^69^ (4.9 nS) and increased it linearly by multiplying it by the free parameter “ChINs inhibitory factor”, in order to explore the possible conductance values that could produce the results obtained *in vivo*. ChINs were fully connected to the population of MSNs.

|  | AMPA | NMDA | GABA |
| --- | --- | --- | --- |
| $\bar{g}_z$ | 6.9nS | 3.45nS | Free parameter |
| $E_z$ | 0mV | 0mV | -70mV |
| $S$ | 0.009 | 0.009 | 1 |
| $\tau_z$ | 6ms | 160ms | 120ms |

#### Dopaminergic modulation

Dopaminergic modulation of dMSNs and iMSNs was implemented following the same equations and parameter values as in^69^ except for the hyperpolarization of dMSNs mediated by the enhancement of the inward-rectifying potassium current (KIR), since this effect was not present in our *in vivo* recordings of spontaneous activity. Briefly, D1 receptor activation produced an enhanced response to depolarising inputs (d←d(1-0.33φ)), while D2 receptor activation produced a small inhibitory effect (k←k(1-0.032 φ)). The parameter φ had an initial value of 0.8, following^69^; we tuned φ in following simulations to reproduce our results obtained *in vivo*. ChINs blockage of activity was implemented as a negative current with a value of 346 following cortical stimulation. Da modulation of cortico-MSN synapses was modified by:

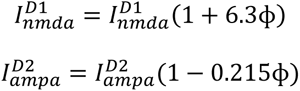

#### Inputs to the model

We provided two different inputs to the model: A spike train following the Poisson process described previously to imitate the sensory stimulation, and a DA switch to reproduce the release of DA during the recordings. The Poisson input consisted on squared pulses of 250ms duration and 5 seconds interstimulus interval. The DA switch was activated randomly in 50% of the stimulations, lasted for 1 second and modified the properties of dMSNs and iMSNs, as well as hyperpolarized ChINs. Each simulation consisted of 20 stimuli.

#### Simulations

During the simulations, two equivalent circuits were simulated together and connected to the same inputs. The only difference consisted in the connectivity of the ChINs to the Poisson generator: In one of the cases, ChINs were connected to evoke visual responses (PT + IT corticostriatal pathway). In the other case, ChINs were not connected to this excitatory input and therefore not triggered by it, mimicking the tactile response in the DMS (IT corticostriatal pathway). Simulations were repeated 10 times and averaged to generate our results.

The model was implemented in Python using the Brian 2 library^136^ and simulated using a fourth-order Runge-Kutta method with a time step of 0.05 ms.

### VIII.#Analysis

#### SWO decomposition using NA-MEMD

Neural oscillations such as the ones recorded in this study are nonlinear^137–139^. Frequency and Time-Frequency analysis are the most common used approaches to analyse the oscillatory properties of neural activity. Nevertheless, traditional decomposition approaches present limitations to analyse this type of data. Therefore, in this study, we used the Noise-assisted Multivariate Empirical Mode Decomposition (NA-MEMD) algorithm^140^ together with Hilbert transform^141^ for the successful analysis of the oscillations of MSNs membrane potential as previously described^15^. Briefly, the original EMD^141^ is a data driven algorithm suitable for nonlinear and non-stationary signals that does not rely on any predetermined template. It decomposes a given signal into a subset of oscillatory modes called Intrinsic Mode Functions (IMFs). Each IMF contains the oscillations of the original data in a certain frequency range, from the fastest to the slowest. Then, Hilbert transform is applied onto each IMF in order to compute its instantaneous frequency and amplitude to retain the original temporal resolution of the signal. The MEMD^140^ is a multivariate extension of the original EMD to n-dimensional signals. The MEMD is computed simultaneously in all dimensions of the signal to ensure the same number of IMFs as output. In addition, new dimensions can be added to the data containing White Gaussian Noise (WGN) to increase its performance, as it has been described that WGN addition reduces mode mixing produced by signal intermittence^142^, acting as a quasidyadic filter that enhances time frequency resolution^141,143^. The application of MEMD to the desired signal together with extra White Gaussian Noise dimensions is known as NA-MEMD analysis^141^. In this study, we applied NA-MEMD algorithm to a multivariate signal composed by the intracellular recording, both LFPs and one extra WGN channel as previously described^15^. The main advantage of this technique is the capacity to deal with the nonlinear and nonstationary properties of neural oscillations^137^ obtaining an enriched decomposition of the oscillatory activity that cannot be achieved by traditional techniques. In order to apply NA-MEMD analysis to our data, we adapted the MEMD Matlab package (http://www.commsp.ee.ic.ac.uk/mandic/research/emd.htm). Standard stopping criterion was described previously^144^. At last, we extracted the IMF carrying the SWO as the one with maximum correlation with the membrane voltage and visually confirmed it in all the recorded cells. Once we isolated the SWO of each recording using NA-MEMD, they were stored for further analysis.

#### Hilbert transform

We computed the frequency of the SWO and theta (6-10 Hz), beta (10-20 Hz) and gamma-bands (20-80 Hz) as the instantaneous frequency using the Hilbert transform^141^. For a given time series x(t), its Hilbert transform H(x)(t) is defined as:

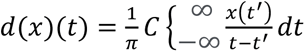

where C indicates the Cauchy principal value. Hilbert transform results in a complex sequence with a real part which is the original data and an imaginary part which is a version of the original data with a 90◦ phase shift; this analytic signal is useful to calculate instantaneous amplitude and frequency; instantaneous amplitude is the amplitude of H(x)(t), instantaneous frequency is the time rate of change of the instantaneous phase angle.

#### Parameters obtained from the evoked sensory responses

In order to quantify the sensory responses (Fig. 3), we first subtracted the IMFs carrying oscillations faster than 50 Hz from the MSNs membrane voltage recordings to eliminate action potentials. In order to extract Up and Down states, we performed a cross-correlation to look for the decomposed IMF that was most similar as possible to the original trace as described in Alegre-Cortés et al., 2021^15^. This IMF contained the SWO. Then, we applied a threshold to that IMF (the mean value plus 0.5 standard deviations) to separate Up from Down states. We developed an interface with custom code written in Matlab to extract the parameters and help us visualize the evoked responses aligned with the trigger. We calculated separately the PSTHs from the evoked visual, tactile or bimodal responses with or without DA during Down and Up states and extracted the mean onset and peak delay, slope, and amplitude for each condition and cell. The onset delay was calculated as the average time between the stimulus trigger and the onset of the evoked potential. The first time derivative of the membrane potential was used to determine the onset of the sensory response within a 250 ms time-window after sensory stimulation. The peak delay was calculated as the average time between the stimulus trigger and the time in which the response reached its maximum peak inside a 250 ms window from the stimulus. The slope was obtained as the first derivative (dv/dt) of the linear fit of the trace between the onset and peak delay time interval, and the response amplitude was defined as the voltage difference between the peak delay and the onset time. The amplitudes of the responses during the Up states were very small or absent, with a rate of failures of 73.95% ± 9.79%, impairing their reliable quantification in most of the cases. Hence, all evoked responses showed in this study are extracted from the Down states, in which at least 14 stimuli were averaged for each condition.

#### Statistical analysis

Wilcoxon Signed rank test was used for comparison of different conditions in matched samples. Wilcoxon Rank-sum test was used for comparison of different conditions in independent samples. The error bars presented in the bar graphs represent the standard error unless stated otherwise. Regarding the boxplots, the red line indicates the median, whereas the bars indicate the lowest and upper 25% of the data values. In the cases in which a linear regression was performed, the values and significance are displayed in the figure legend. Values exhibited in the tables represent the mean ± standard deviation. When required, alpha values for multiple comparisons were corrected using Holm-Bonferroni correction. Confidence level was set to p=0.05. All statistical analyses were done in Matlab (Mathworks).

## SUPPLEMENTARY INFORMATION

**Supplementary Figure 1.**
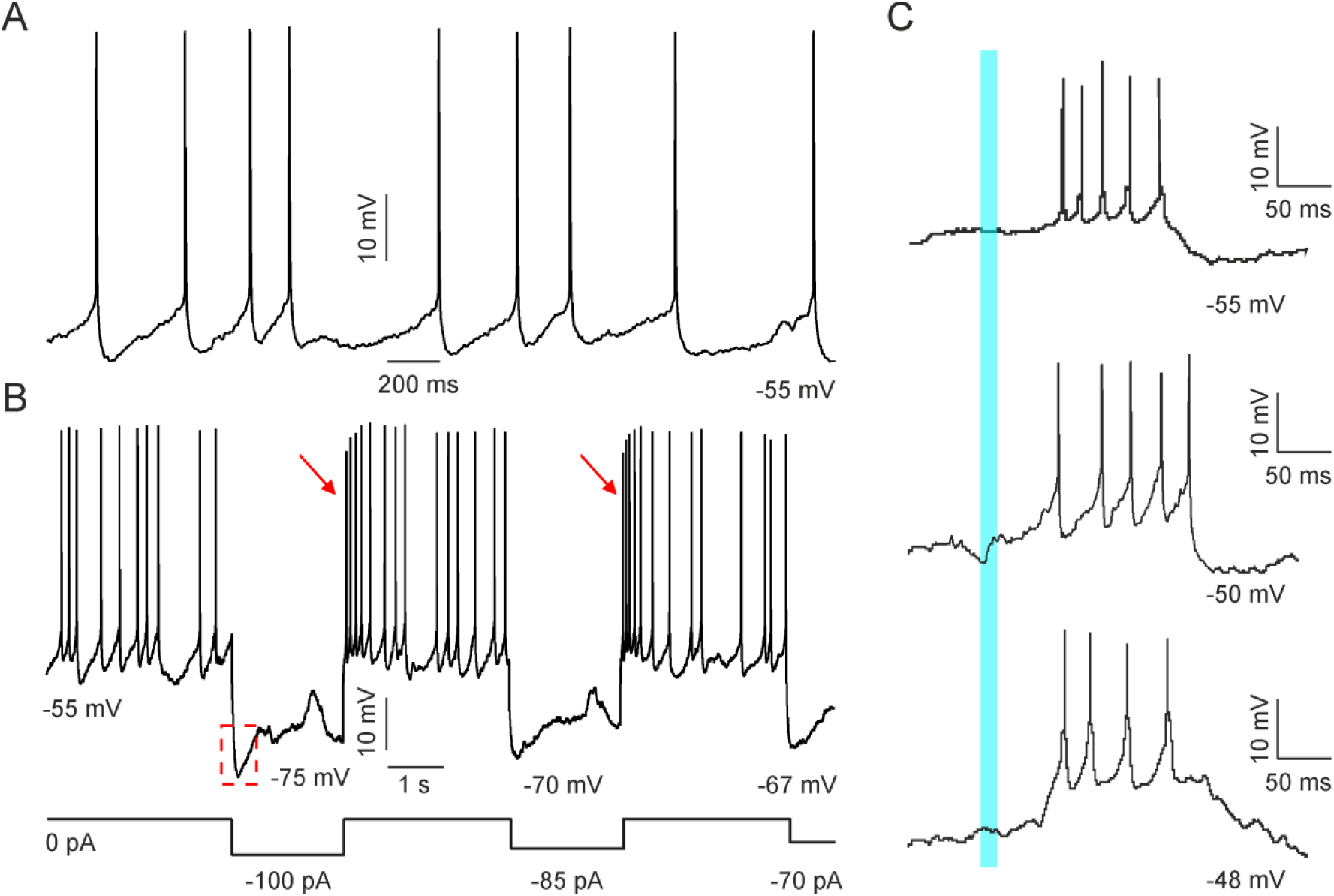
*In vivo* cholinergic interneuron recording. **A,** Example of the spontaneous activity of a cholinergic interneuron recorded *in vivo*. Notice its spontaneous tonic activity. **B,** Response of the same ChIN to step current injections. Notice the voltage sag response (red dashed trace) and the rebound spikes characteristic of cholinergic interneurons (red arrows). **C.** Trace example of 3 DMS-ChINs responding to visual stimulation. Blue line indicates the sensory stimulation.

**Supplementary Figure 2.**
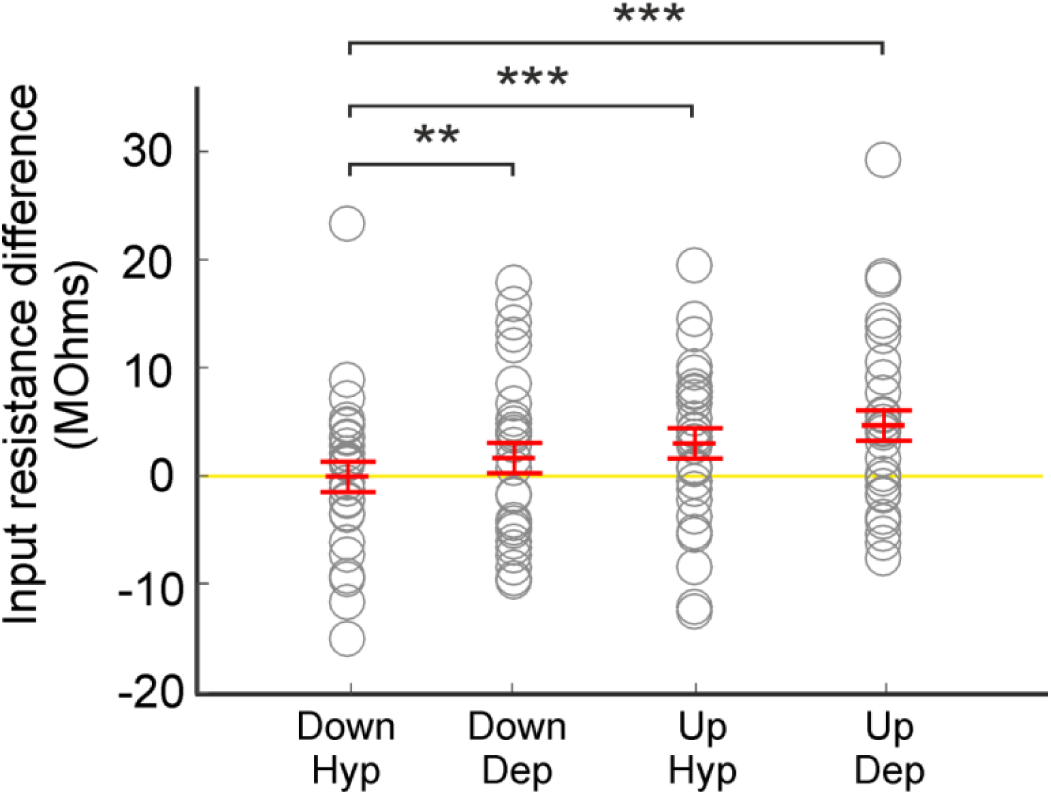
MSNs input resistance. Differences in the input resistance of MSNs during Up and Down states calculated during negative and positive current steps. All values are subtracted with respect to the “Down Hyp” condition. Yellow line indicates the value 0. n=33. **p<0.01; *** p<0.001.

**Supplementary Figure 3.**
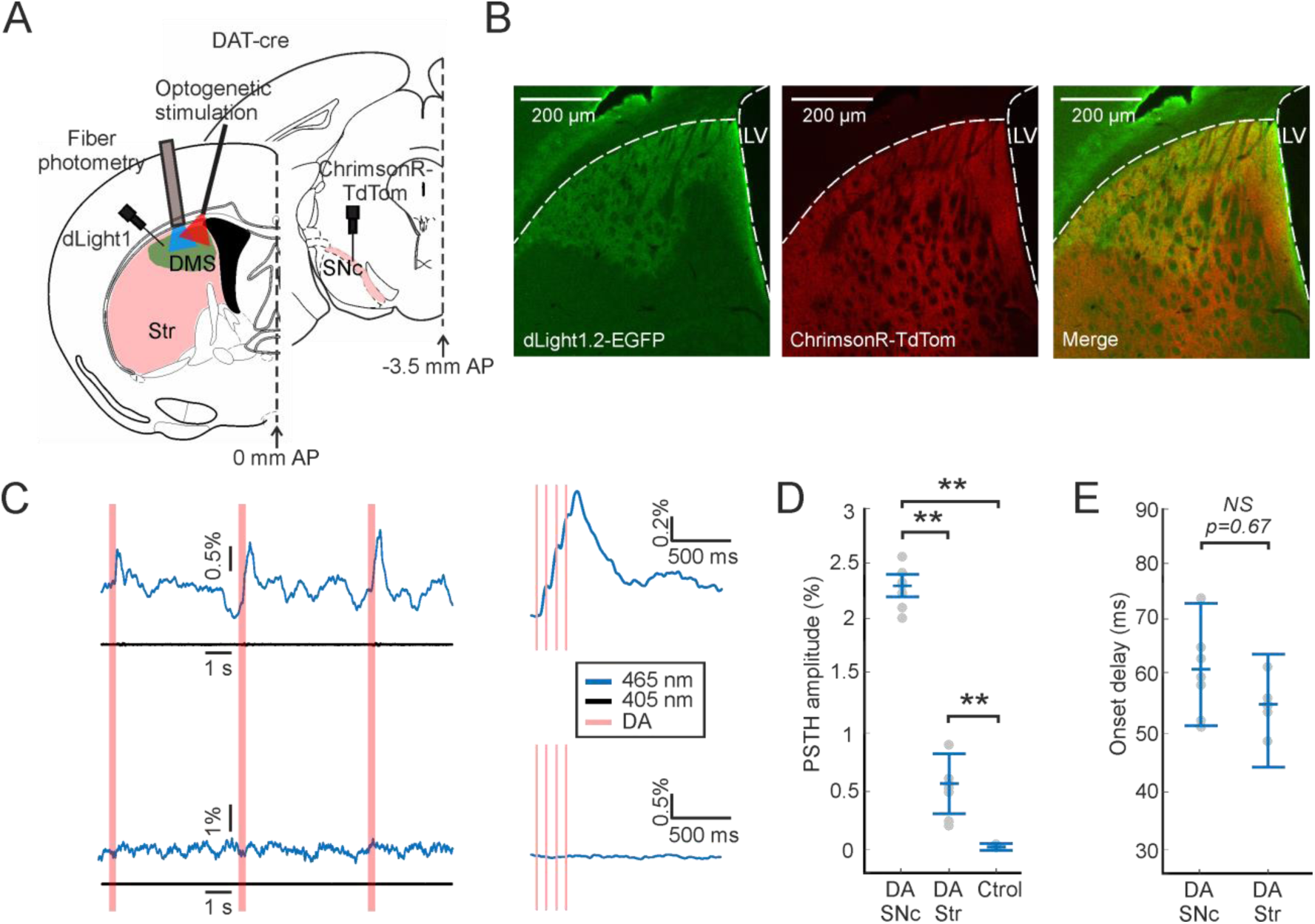
Optogenetic dopamine release in the DMS. **A,** Schematic representation of the fiber photometry experimental set up. **B,** Coronal images of a DAT-cre mice injected with the viruses enconding the DA sensor dLight1-EGFP and the opsin ChrimsonR-TdTomato in the DMS and SNc respectively. Left: Image showing the expression of the dLight1 sensor injected in the dorsal striatum. Middle: Image showing the expression of the ChrimsonR opsin in the dopaminergic terminals. Right: Merge of both images. **C,** Left: Trace example of the fluorescence measurements along time from one of the recording sessions. Right: Waveform average of the elicited DA release when optogenetically stimulating the dopaminergic terminals in the DMS infected with the virus expressing the opsin ChrimsonR (top) or with the control virus (bottom). Blue traces correspond to the 465 nm signal. Black traces correspond to the isosbestic 405 nm signal (used as a reference point) [see methods]. Red bars represent the time of the optogenetic stimulation trains (peak excitation wavelength of ∼660 nm). **D,** Quantification of the changes in fluorescence when stimulating directly the SNc (DA SNc), the dopaminergic terminals in the DMS (DA Str) or during control conditions (Ctrol). ** p<0.01. **E,** Quantification of the onset delays from the fluorescence changes when stimulating directly the SNc (DA SNc) or in the dopaminergic terminals in the DMS (DA Str).

**Supplementary Figure 4.**
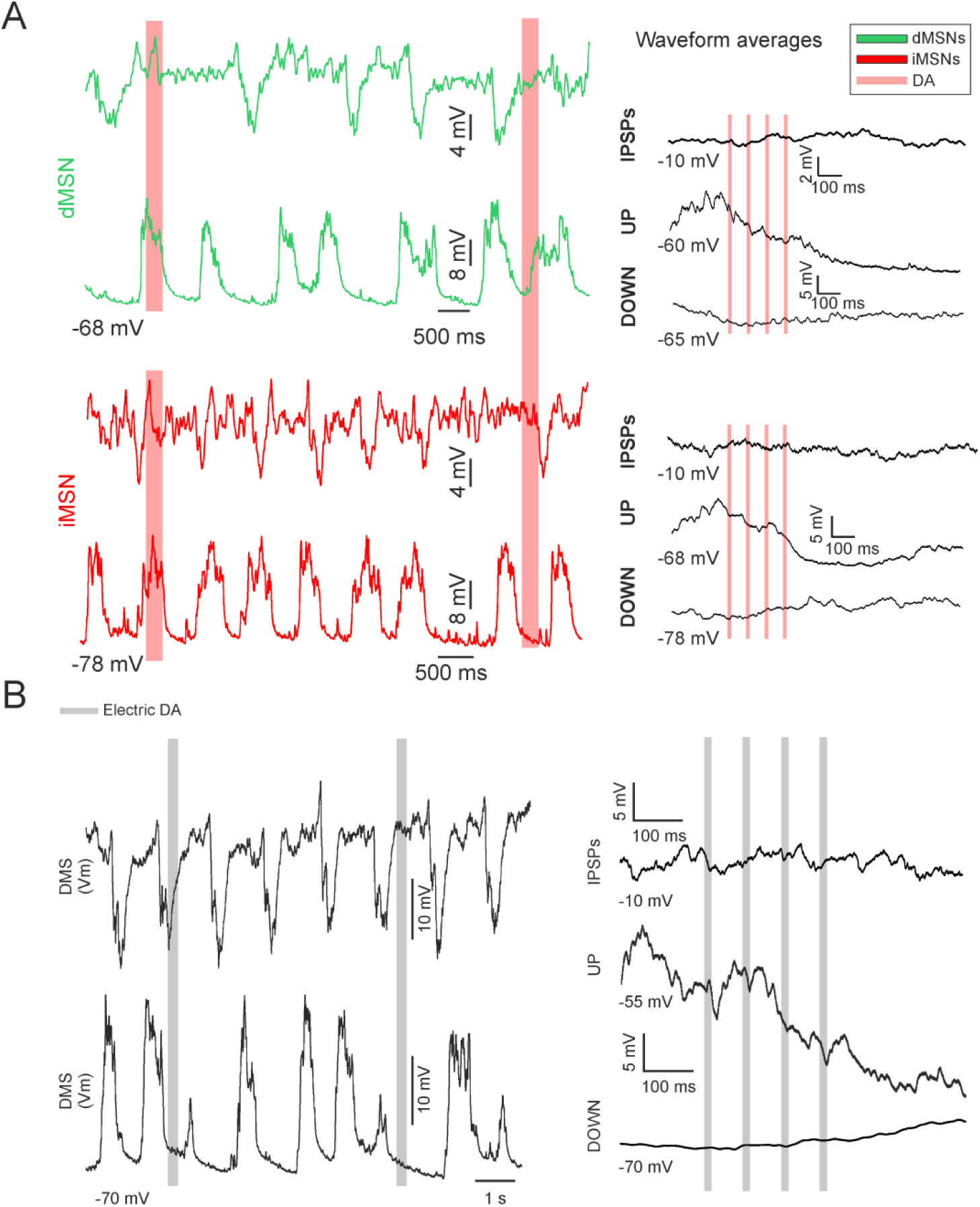
Electric stimulation of SNc dopaminergic neurons. **A,** left: Example trace of an in vivo whole-cell recording of a DMS dMSNs (green) and iMSNs (red) dorsomedial MSN while stimulating DMS dopaminergic terminals optogenetically every 5 seconds. Right: Waveform averages of the same MSNs at different membrane potentials. Coloured lines represent optogenetic stimulation. **B,** Left: Example trace of an *in vivo* whole-cell recording of a dorsomedial MSN while stimulating electrically in the SNc dopaminergic neurons every 5 seconds. Upper trace corresponds to the same neuron while depolarizing it to see the inhibitory components. Right: Waveform averages of the same MSN at different membrane potentials. Coloured lines represent electric stimulation.

**Supplementary Figure 5.**
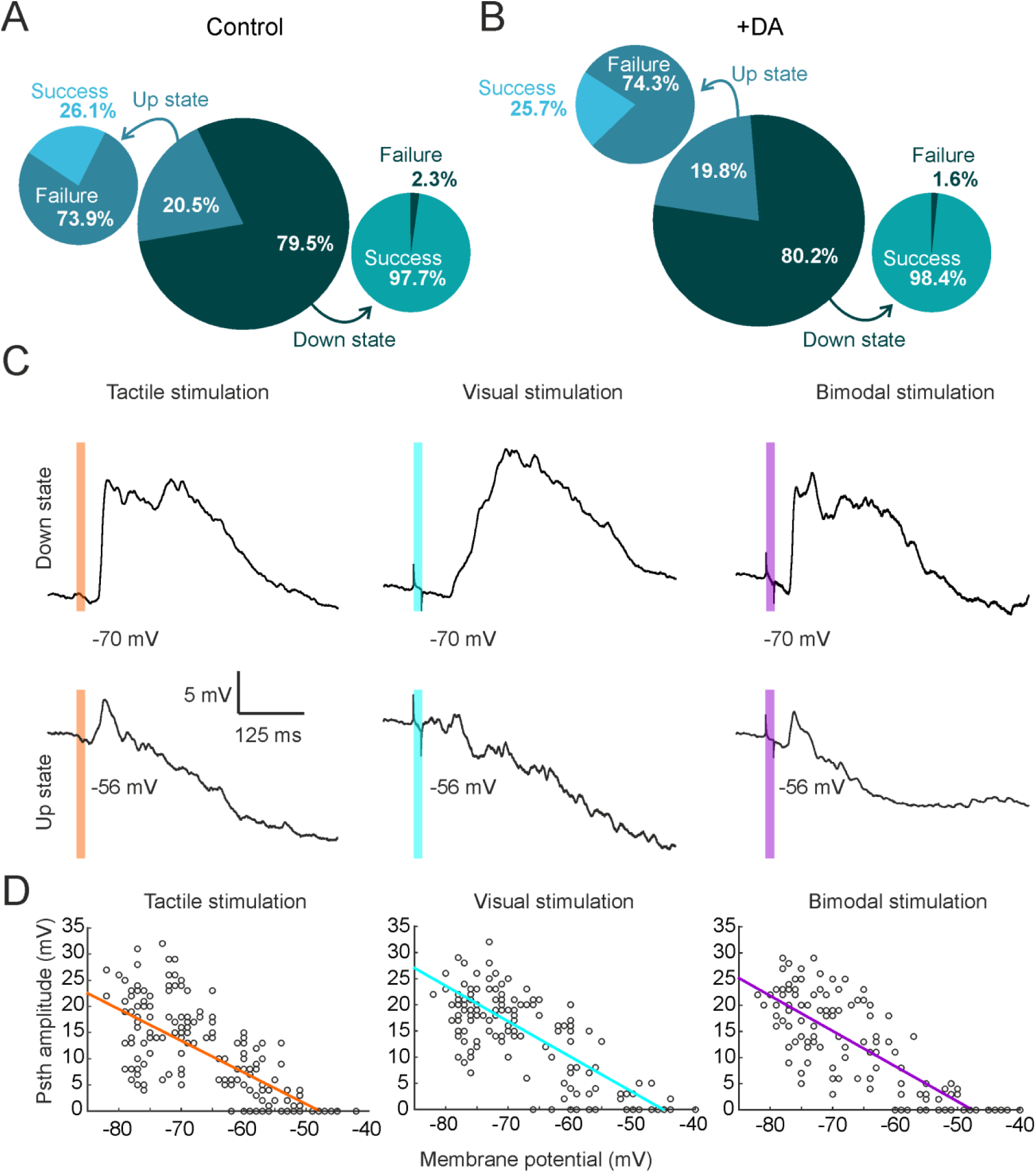
Voltage-dependence of sensory evoked responses. A, B, Quantification of the probability that a sensory stimulus happens in the Up (light colour) or Down state (dark colour) (big circle), and the probability of failure (dark colour) or success (light colour) of that stimuli to evoke a sensory response (small circles) with (A) or without dopamine release (B). **C,** Waveform averages of an example MSN responding to tactile (left), visual (middle) and bimodal (right) stimulation during Down (upper traces) or Up states (bottom traces). Total trials in Down state/cell n=51. Total trials in Up state/cell n=11. The coloured line represents the sensory stimulation. **D,** Variation of amplitudes at different membrane potentials corresponding to Up and Down states for tactile (left), visual (middle) and bimodal (right) stimulation. For each cell, a representative sample of amplitudes at different membrane potentials was used. Each point represents each one of these samples. A linear regression between -85 and -40 mV values (Tactile: R=0.78, R^2^=0.61, p<0.0001; Visual: R=0.82, R^2^=0.68, p<0.0001; Bimodal: R=0.81, R^2^=0.67, p<0.0001) illustrates the voltage dependence of the sensory responses.

**Supplementary Figure 6.**
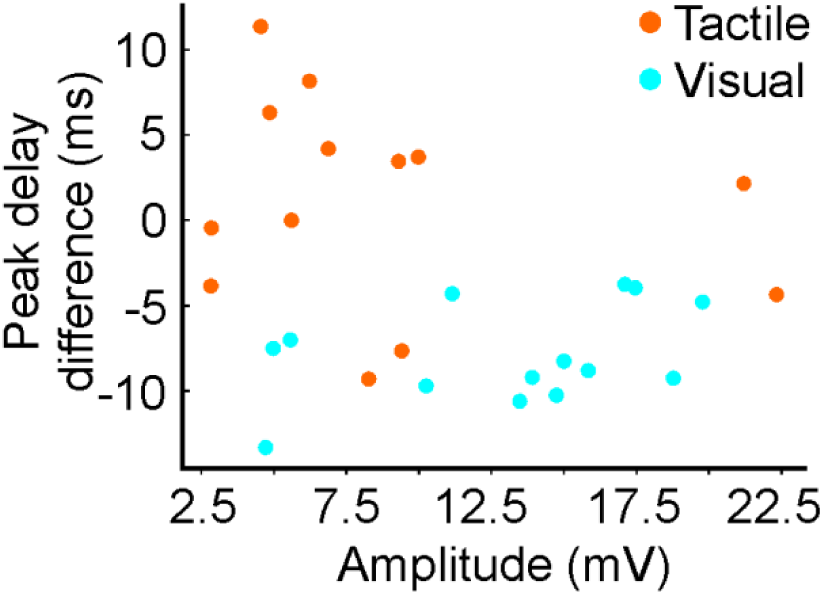
No dependence between the peak delay difference and amplitude of tactile and visual responses. Peak delay differences (DA – control condition) for visual and tactile responses comparing them with their respective control amplitudes in dMSNs. Each point represents a cell. No correlation was found when performing a linear regression (Tactile: R=-0.22, R^2^=0.04, p=0.46; Visual: R=0.33, R^2^=0.10, p=0.24).

**Supplementary Figure 7.**
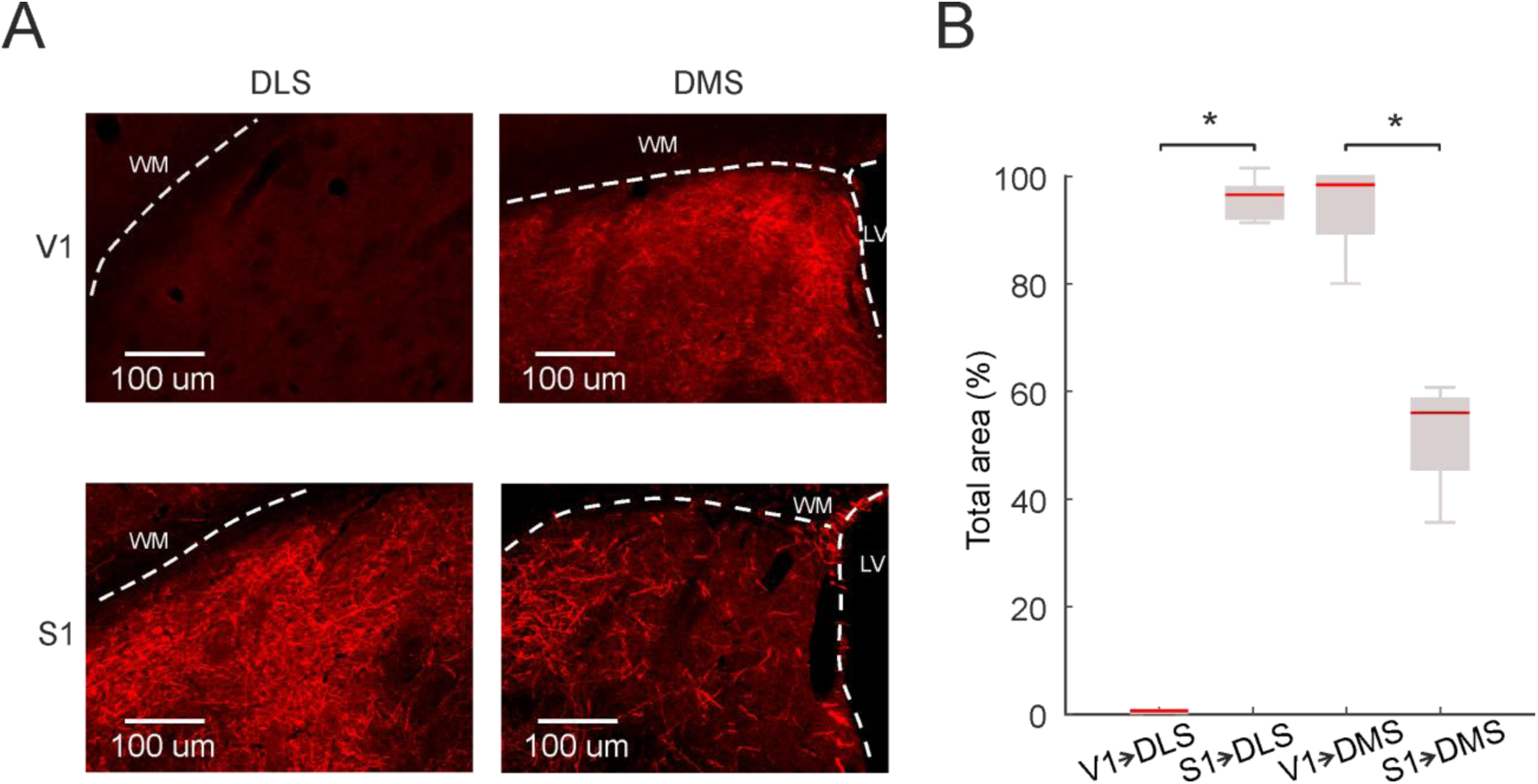
V1 and S1 projections towards the DLS and DMS. **A,** Representative image of BDA labelled projections from V1 (top) and S1 (bottom) towards the DLS (left) and the DMS (right). **B,** Quantification of the covered area by the labelled axons from V1 and S1 towards the DLS and DMS.

**Supplementary Table 1.** Mean values of tactile and visual evoked responses in S1 and V1 LFP recordings with or without DA release. n= 12. All values are mean ± standard deviation.

| | Onset delay<br>(ms) | Peak delay<br>(ms) | Amplitude<br>(μV) | Slope<br>(μV/ms) |
| --- | --- | --- | --- | --- |
| Tactile responses evoked in S1 | $12.75 \pm 3.91$ | $40.25 \pm 5.12$ | $521 \pm 42.42$ | $18.94 \pm 5.52$ |
| Tactile responses evoked in S1 + DA | $11.65 \pm 3.73$ | $43.19 \pm 6.17$ | $540 \pm 40.69$ | $17.13 \pm 4.28$ |
| Visual responses evoked in V1 | $49.23 \pm 10.87$ | $95.26 \pm 18.49$ | $505 \pm 45.41$ | $11.67 \pm 4.59$ |
| Visual responses evoked in V1 + DA | $50.74 \pm 11.97$ | $97.65 \pm 21.46$ | $499 \pm 50.23$ | $10.21 \pm 5.13$ |

**Supplementary Table 2.** Description of the animals used in the electrophysiological recordings. The first three rows are total values, the rest show mean ± standard deviation. n=15 (12 + 3 control).

|  |  |
| --- | --- |
| Animals | 15 |
| Males | 9 |
| Females | 6 |
| Weight (gr) | $33.33 \pm 10.42$ |
| Male weight (gr) | $38.66 \pm 9.82$ |
| Female weight (gr) | $25.33 \pm 4.67$ |
| Age (weeks) | $22.4 \pm 8.55$ |
| Male age (weeks) | $24.66 \pm 10.09$ |
| Female age (weeks) | $19 \pm 4.33$ |
| Cells/animal | $2.86 \pm 1.09$ |
| Cells/male | $2.78 \pm 1.13$ |
| Cells/female | $3.16 \pm 0.98$ |

## Notes

### Competing Interest Statement

The authors have declared no competing interest.

